# Expansion of mammary intraepithelial lymphocytes and intestinal inputs shape T cell dynamics in lactogenesis

**DOI:** 10.1101/2024.07.09.602739

**Authors:** Abigail Jaquish, Eleni Phung, Xutong Gong, Pilar Baldominos, Silvia Galvan-Pena, Isabelle Bursulaya, Ian Magill, Eleonara Marina, ImmgenT consortium, Kerri Bertrand, Christina Chambers, Andrés R. Muñoz-Rojas, Judith Agudo, Diane Mathis, Christophe Benoist, Deepshika Ramanan

## Abstract

Pregnancy brings about profound changes in the mammary gland to prepare for lactation, yet immunocyte changes that accompany this rapid remodeling are incompletely understood. We comprehensively analyzed mammary T cells, revealing a marked increase in CD4+ and CD8+ T effector cells, including an expansion of TCRαβ+CD8αα+ cells, in pregnancy and lactation. T cells were localized in the mammary epithelium, resembling intraepithelial lymphocytes (IELs) typically found in mucosal tissues. Similarity to mucosal tissues was substantiated by demonstrating partial dependence on microbial cues, T cell migration from the intestine to the mammary gland in late pregnancy, and shared TCR clonotypes between intestinal and mammary tissues, including intriguing public TCR families. Putative counterparts of mammary IELs were found in human breast and milk. Mammary T cells are thus poised to manage the transition from a non-mucosal tissue to a mucosal barrier during lactogenesis.

## INTRODUCTION

The mammary gland is remarkable in its capacity to undergo multiple cycles of growth and regression during reproductive years. Mammary gland remodeling is largely driven by hormonal cues, resulting in stage specific tissue adaptations during puberty, pregnancy, lactation, and involution^1, 2, 3^. Pregnancy initiates proliferation of mammary epithelial cells and ductal branching to support lactogenesis, a crucial process that ensures successful production and transfer of breast milk, essential for offspring health^4, 5^. The transition of the mammary gland into a secretory organ for the duration of lactation involves dramatic restructuring of cell composition and tissue enlargement, and increases its exposure to the outside environment, including microbes on maternal skin and offspring’s oral cavity, rendering it a temporary barrier tissue^6, 7^. The end of lactation triggers mammary involution, a reversal to its non-secretory state marked by extensive apoptosis and tissue shrinkage^8, 9^. The physiological stress that accompanies the rapid transformation of the mammary gland requires an extensive support network, including immunocytes, but the types of immunocytes involved and their functions in lactogenesis are unclear.

Innate and adaptive immunocytes are involved in immune regulation of the mammary gland at various developmental stages. Mast cells and eosinophils promote ductal branching during mammary gland development in puberty^10, 11^. Macrophages are required for mammary gland morphogenesis in puberty, alveologenesis in pregnancy, and tissue regression in involution^11, 12, 13, 14, 15, 16^. A specialized population of macrophages called ductal macrophages dominate the lactating mammary gland, and another subtype of macrophages, lactation-induced macrophages (liMacs), have been recently identified to support lactogenesis and milk production^12, 17^. B cells, specifically IgA- and IgG-producing plasma cells, are abundant in lactation and promote offspring health by shaping the antibody composition of milk. We and others have shown that IgA-producing plasma cells can migrate from the intestine to the mammary gland in a CCL28 dependent manner^18, 19, 20, 21, 22^. Despite recent advances, our comprehension of various immunocyte types and their collaborative roles in facilitating mammary remodeling and regulating milk composition remains limited.

T cells play a fundamental role in tissue homeostasis by promoting defense, tolerance, tissue repair, and regulating cell proliferation and death. CD4+ T cells have been described in the mammary gland in puberty, and a subset of CD4+ regulatory T cells (Tregs) that express RORγ increase during involution^23, 24^. In addition, CD4+ and CD8+ T cells are present in milk^25^. Mammary ψο+ T cells with innate-like properties increase during lactation, and lack of these ψο+ T cells can favor mammary oncogenesis^26^. During lactogenesis, several processes occur simultaneously, including epithelial cell stress from rapid expansion and milk production, and heightened exposure to the sudden increase in newly revealed self-antigens, which could create a conundrum for T cell tolerance, highlighting the need for T cell assistance. Yet, there is a large gap in our understanding of which T cell subsets are involved in lactogenesis, and how they contribute to tissue specific adaptations in the mammary gland during gestation, lactation, and involution.

We set out to define immunocyte changes in lactogenesis and uncovered novel T cell dynamics in the mammary gland. We provide a detailed overview of mammary intraepithelial T cells, shaped by intestinal and microbial influences, that accompany the remodeling of the mammary gland into a mucosal-like state during lactogenesis.

## RESULTS

### Lactation leads to increased T cell activation in the mammary gland

The size and composition of the mammary gland undergo substantial changes in preparation for lactation, yet there remains a gap in our understanding of how immunocytes adapt to, and perhaps facilitate, those transitions. We first quantified mammary immunocytes (CD45+) across developmental stages by flow cytometry and found that gestation (G) initiated a rapid increase in the total number of immunocytes, that was maintained in lactation (L), and involution (I) (Fig. 1A). To determine the expanding cell types and chart their adaptations, we performed a temporal analysis of immunocytes across different stages in female C57BL/6 (B6) mice; profiling by single-cell RNA sequencing (scRNAseq) mammary glands from nulliparous, gestation (day 17), early lactation (day 3 postpartum), and involution (day 1 post-weaning) stages, multiplexed into the same runs. A total of 60,060 immunocytes were captured across all lineages from 20 mice across three independent runs, revealing increased representation of T cell populations (Fig. S1A-B). Validation by flow cytometry confirmed the expansion of T cells, while the proportion of myeloid cells, which dominated in the nulliparous, declined somewhat (while remaining the largest cell populations, as previously described^12, 17, 24^)(Fig. 1B). Analysis of the scRNAseq data revealed several myeloid cell populations, including the recently described lactation associated liMacs, specifically during lactation^17^ (Fig. S1B,C). While we did not observe shifts in B cell populations, we identified CD103+ NK cells that were associated with lactation (Fig. S1A, D). To better understand the expanding T cell populations, we applied Louvain clustering^27^, which parsed 6 clusters of T cells, which were annotated as naïve T cells (Tn), and effector T cells (Teff) that were either CD4+, CD8αβ+ or CD8α+CD8β-(CD8αα), and CD4-CD8-CD3+ (Double negative, DN) (Fig. 1C, replicate in Fig. S2A) (gene signatures from ImmGen^28, 29, 30^). There was a strong shift in the relative proportions of naïve vs activated states of CD4+ T cells and CD8+ T cells, from mostly naïve T cells before and during gestation to mostly activated states during lactation, with a particularly striking increase in CD8αα+ Teff in late pregnancy that persisted during involution (Fig. 1C). This shift was observed among all TCRβ+ populations in the mammary gland by flow cytometry (Fig. S2B).

**Figure 1.**
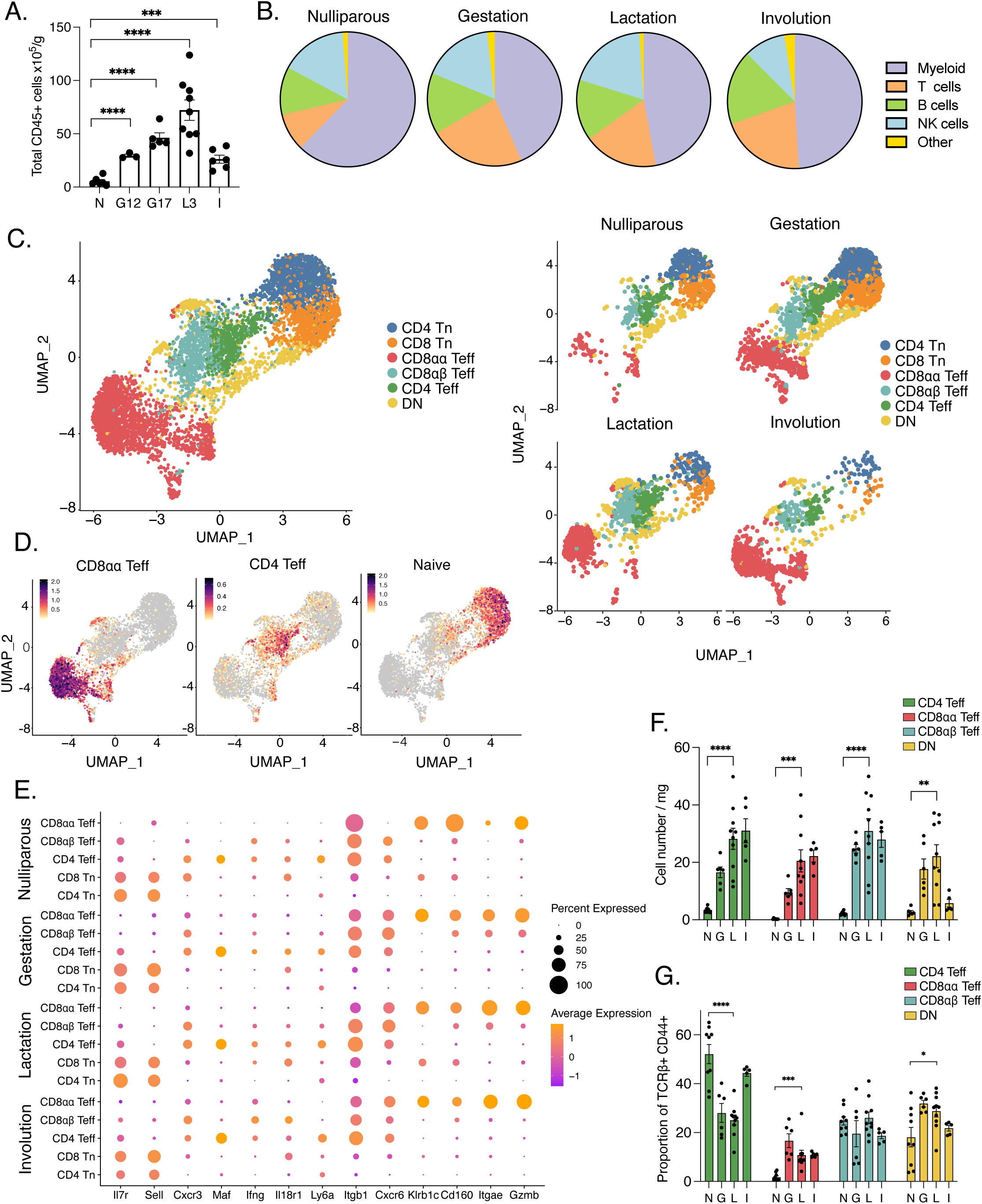
Late gestation and lactation lead to increased Teff populations in the mammary gland. A) Quantification of total number of CD45+ cells normalized to mammary gland weight across stages of gestation and lactation by flow cytometry. N=nulliparous (n=6), G12=gestation day 12 (n=3), G17=gestation day 17 (n=5), L3=lactation days 3-5 (n=9), and I=involution, 1 day post weaning (n=6). B) Representative proportions of major immune cell types in the mammary gland across stages, nulliparous (n=5), gestation (n=3), lactation (n=5), and involution (n=5), quantified by flow cytometry. C) UMAP projection of mammary T cells. Split by stages: nulliparous, gestation (G17), lactation (L3), and involution (right). Representative UMAP is from one of three independent experiments, n=3. D) Feature plots of CD8αα Teff, CD4 Teff and Naïve T cells from (C). E) Dot plot of selected highly expressed genes in T cell clusters across stages identified in (C). Dot size represents the percentage of cells expressing the selected gene and color indicates expression level. F-G) Quantification by flow cytometry of cell numbers (F) and proportions (G) of T cell populations identified in (C) normalized to mammary gland weight. Teff populations were determined as CD4 Teff: CD4+CD44+CD62L-. CD8αα Teff: CD8α+CD8β-CD44+CD62L-. CD8αβ Teff: CD8α+CD8β+CD44+CD62L-. DN: TCRβ+CD4-CD8α-. N=nulliparous (n=8, n=6 for DN), G=gestation (n=6), L=lactation (n=10) and I=involution (n=5). **p<0.01, ***p<0.001, ****p<0.0001 by two tailed student’s t-test (A, F, and G). Data is representative of ≥3 independent experiments, bars in plots indicate mean ± SEM.

In contrast to the CD8αβ+ Teff cells characterized by *Itgb1* and *Cxcr6* expression, CD8αα+ Teff cells also expressed high levels of *Klrb1c*, *Cd160, Itgae*, and *Gzmb*, suggesting increased cytotoxic potential and tissue residency (Fig. 1E, S3). CD8αα+ Teff cells also expressed increased cell adhesion and proliferation genes such as *Mcm5, Mcm7, Mki67, Lgals1*, and *Hmmr* in differential levels across stages (Fig. S2C-D, S3). The transcriptional signature of CD8αα+ cells was reminiscent of CD8αα+ T cells^28, 30^ that have innate properties and reside in the epithelium of mucosal tissues, referred to as intraepithelial lymphocytes (IELs) (intestinal CD8αα+ IEL signature^31,30,31^ applied to mammary CD8αα+ cluster in Fig. 1C). A similar signature has also been described in TCRαβ+ innate-like T cells with high cytotoxic potential in mammary tumors called αβILTCKs^32^ (Fig. S2E). CD8αα+ T cells mostly differentiate in the thymus in response to strong agonists^32, 33, 34, 35, 36^ and use either TCRαβ or TCRψο, but mammary CD8αα+ T cells were mostly TCRαβ+ (Fig. S4A). Flow cytometric validation confirmed that gestating and lactating mammary glands displayed a significant increase in the proportion of CD8αα+ and DN T cells and in cell numbers of CD4+ Teff, CD8+ Teff, and DN populations (denoted by CD44+ staining) (Fig. 1F-G, gating strategy in S4B).

We investigated other T cell populations and interestingly, while Tregs increased during gestation, lactation led to a significant drop in Treg proportions and numbers (Fig. S4C), followed by an increase in involution. Unconventional T cell subsets such as ψο T cells, MR1+ MAIT cells and iNKT cells were sparse, and to some extent, decreased upon lactation (Fig. S4D,E). Thus, mammary gland remodeling is accompanied by strong changes in T cell populations, including a hyper-activated state and increased CD4+ Teff, CD8αβ+ Teff, and CD8αα+ Teff cells in lactation, presumably in response to rapid epithelial cell proliferation and exposure to the outside environment.

### Mammary T cell populations are located in the epithelium

The expansion of mammary CD8αα+ T cells, an abundant cell-type in the intestine, was intriguing. To compare transcriptional similarities of CD8αα+ T cells across tissues, we multiplexed nulliparous and lactating mammary gland, small and large intestine, and spleen from the same mouse into scRNAseq experiments (Fig. 2A, 25,096 cells, 4 mice, 2 independent experiments). At first glance T cells from the same tissue clustered together suggesting a distinct transcriptional state that was tissue specific (Fig. 2A). However, signatures of effector CD4+, CD8αβ+, and CD8αα+ T cells were similar across organs (Fig. 2B). Most CD8αα+ T cell genes, *Tyrobp*, *Fcer1g*, *Itgae*, *Gzmb*, displayed similar expression in mammary gland, small intestine and large intestine but not the spleen (Fig. S5A). The expression of killer cell lectin-like receptor (KLR) family genes such as *Klra1, Klra7, Klrb1a,* and *Klrb1c*, were higher in mammary CD8αα+ T cells compared to intestinal tissues (Fig. S5A). We validated two of the classical IEL markers used above by flow cytometry, Ly49a (*Klra1*) – a killer cell lectin like receptor that binds MHC-I, and CD103 (*Itgae*), an integrin that mediates cell adhesion and tissue retention by binding to e-cadherin on epithelial cells. In line with the gene expression data above, lactation led to increased Ly49a expression in CD103+ CD8αα+ T cells in the mammary gland and to a lesser extent in the small intestine (Fig. 2C, S5B). Lactation also led to a decrease in CD103+ CD8αα+ T cells in the small intestine but not the mammary gland (Fig. 2C-D, S5B). We also observed a lactation-mediated increase in the expression of markers associated with intestinal IELs, such as CD160, CD38, and 2B4 (CD244)^37, 38, 39^, on mammary CD8αα+ and CD8αβ+T cells (Fig. 2D, S5C). Based on the gene signatures and surface expression profiles, we hypothesized that CD8αα+ T cells in the mammary gland could be IELs. Since the defining characteristic of IELs is their residence in the epithelial layer, we surveyed the physical location of the effector T cells in the mammary gland tissue. Indeed, we observed CD8αα+ T cells adjacent to both basal and luminal epithelial cells (basal epithelial cells by Krt14 and luminal epithelial cells by Krt8), in increased numbers during gestation and early lactation (Fig. 2E and S5D). Surprisingly, CD4+ and CD8αβ+ T cells were also intraepithelial in location (Fig. 2E and S5D). Overall, our results demonstrate that mammary CD8αα+ T cells have marked phenotypic similarity to intestinal CD8αα+ IELs. Mammary CD4+ and CD8αβ+ T cell populations are also IELs, reminiscent of mucosal epithelium. The increase in mammary IELs in lactation is indicative of a temporary mucosal state of the reconfigured mammary gland.

**Figure 2.**
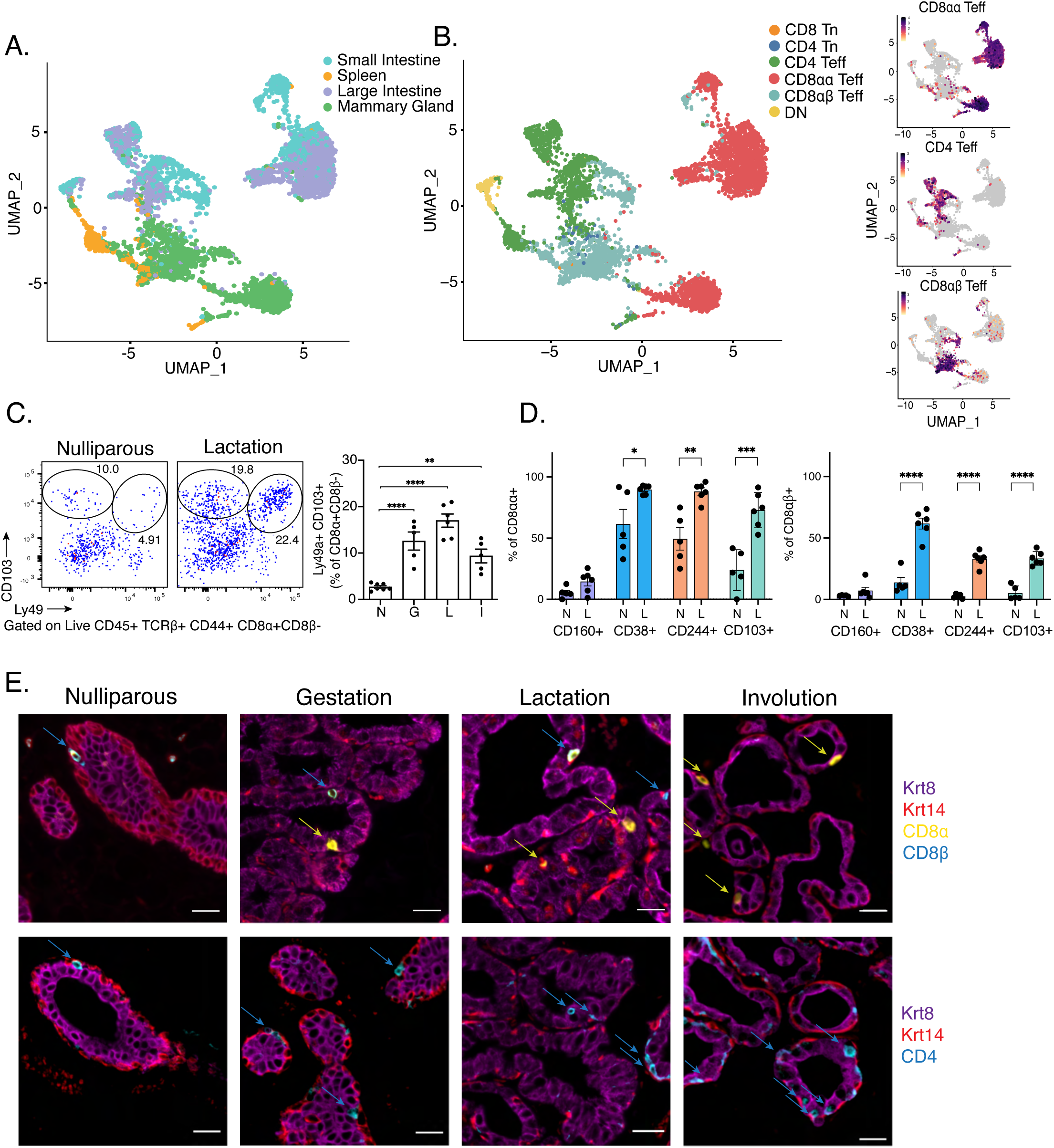
Mammary T cells are intraepithelial lymphocytes. A) Summary UMAP projections of single cell sequencing performed on T cells from mammary gland, large intestine, spleen and small intestine of lactating mice. B) UMAP and feature plots showing the transcriptional localization of featured T cell signatures. C) Representative flow cytometry plots and quantification of CD103+Ly49+CD8αα+ T cells (gated on Live CD45+TCRβ+CD8⍺+CD8β-) across gestational and lactation stages in the mammary gland. N=nulliparous (n=7), G=gestation day 17 (n=5), L=lactation days 3-5 (n=6) and I=involution,1 day post weaning (n=5). D) Proportion of CD8αα+ (left) and CD8αβ+ (right) cells that express CD160, CD38, CD244, and CD103 in nulliparous (n=5) and lactation (n=6) mammary glands. E) Representative immunofluorescence images of the mammary gland at nulliparous, gestation (G17), lactation (L3), and involution. Epithelial cells (Krt8+ luminal cells in magenta and Krt14+ basal cells in red), and T cells, CD8⍺ in yellow, CD8β in cyan (top) and CD4 in cyan (bottom). Scale bar = 20µm. *p<0.05, **p<0.01, ***p<0.001, ****p<0.0001 by two tailed student’s t-test (D). Representative of ≥3 independent experiments, bars in plots indicate mean ± SEM.

### Putative mammary T cell-epithelial cell interaction networks shift during lactation

Given the intraepithelial location of expanding mammary T cells during lactation, we investigated the potential interaction pathways between mammary IELs and basal and luminal epithelial cells in nulliparous and lactating mammary glands using CellChat^40^ (25,506 cells, 4 mice per condition). [CellChat uses a manually curated database (CellChatDB) of literature-supported ligand:receptor signaling pathways, including multisubunit structures, cofactors, coreceptors, agonists and antagonists. Each potential interaction is assigned an interaction probability score based on the law of mass action to model the likelihood of an interaction based on the expression of the ligand, receptor, and any cofactors. Statistically significant interactions are identified through a permutation test on randomly assigned group labels for cells]. We first identified differentially expressed genes (p < 0.05) between nulliparous and lactating mice for each cell population, and then mapped their projected interactions based on the fold change of ligands and receptors. For visualization purposes, we combined ligand:receptor pairs into functionally related signaling pathways (Fig. 3A,D, S6A,C), and plotted communication probabilities between ligand:receptor pairs upregulated (Fig. 3B, S7) or downregulated (Fig. S6B,D) with lactation. Potential interactions that were upregulated with lactation were enriched in pathways related to cell adhesion and migration, including *Pecam1,* selectins (*Sell)*, laminins (*Lamb3*), and galectins (*Lgals9*) (Fig. 3A,B). Expression of *Lgals9* and *Lamb3* transcripts was increased in CD8αα+ and CD8αβ+ T cells, while *Pecam1* and *Sell* expression was increased in CD4+ Teff cells and DN T cells (Fig 3C). Interestingly, *Sell* was highly expressed in nulliparous CD8αβ+ Teff cells, but its potential interacting partner shifted from *Podxl* in nulliparous mice to *Glycam1* in lactation (Fig. 3B, S6B). *Glycam1* is a mucin-like glycoprotein produced by luminal cells in a prolactin-dependent manner^41^, and could potentially facilitate epithelial-IEL interactions during lactogenesis.

**Figure 3.**
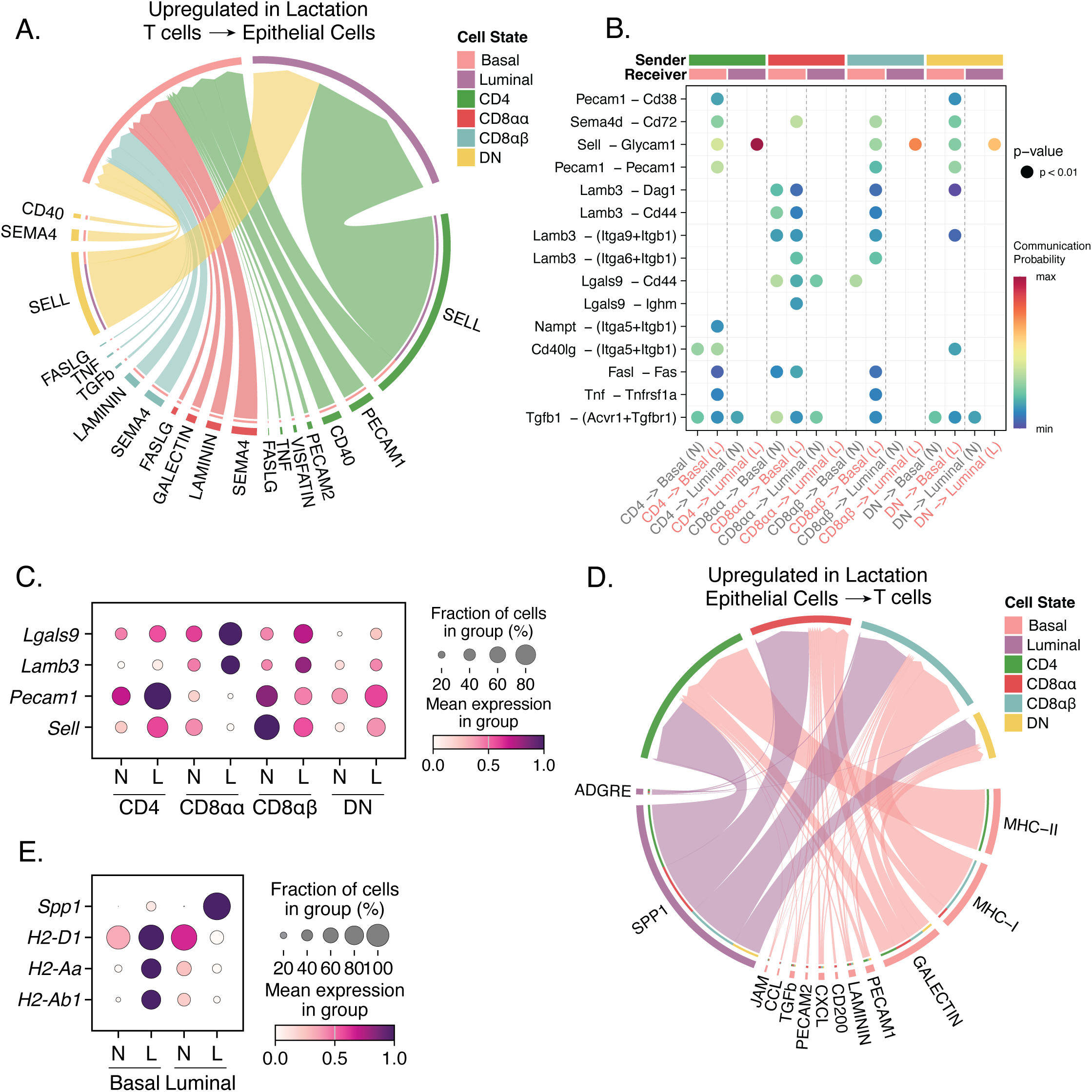
Putative interaction networks between mammary T cells and epithelial cells. A, D) Chord diagrams showing potential signaling pathways upregulated in lactation from T cell populations to epithelial cells (A) and from epithelial cells to T cell populations (D). Ligand:receptor pairs as summarized into functionally related signaling pathways. Outer thicker bars represent the cell population that is the source or target of the signaling pathway in the chord diagram. The inner thinner bar color is the target of the signal. The thickness of the edge represents the signaling strength (communication probability) as calculated by CellChat. B) Dot plot showing the communication probabilities of ligand:receptor pairs upregulated in lactation from T cells to epithelial cells. Heatmap depicts the communication probability of each ligand pair for each cell pair in nulliparous (N) and lactating (L) mammary glands. Sender and receivers are indicated by the color bars on top. C, E) Dot plot of transcript expression levels in mammary IELs (C) and epithelial cells (E) in nulliparous (N) and lactating (L) mammary glands, depicting the percent of cells expressing and mean expression levels in each cell population.

Predicted interactions from epithelial cells to T cells were enriched for immunoregulatory pathways in lactation, including increased expression of MHC complexes in basal epithelial cells with MHC I signaling to CD8αα+ and CD8aβ+ IELs, and MHC II signaling to CD4+ Teff cells (Fig. 3D,E, S7). Another strongly predicted interaction between luminal cells and multiple IEL populations involves osteopontin (*Spp1*), a glycoprotein associated with epithelial cell proliferation and local immunity during lactation^42^ (Fig. 3D,E and S7). Thus, putative interaction analysis suggests multiple signaling pathways between epithelial cells and intraepithelial T cells which could regulate immune surveillance and lactogenesis, providing candidates for future functional studies.

### T cells migrate from the intestine during gestation

Intestinal TCRαβ+ CD8αα+ IELs arise from thymic progenitors acquiring their effector program and expression of gut-homing receptors in the thymus as a result of agonist stimulation by self-antigens ^32, 33, 34, 35, 36^. To test whether the strong increase in mammary gland CD8αα+ IELs stemmed directly from the thymus, we thymectomized female mice before pregnancy (4 weeks of age) and assessed mammary gland CD8αα+ IELs during lactation. Surprisingly, there were no differences between thymectomized and control mice in numbers of CD8αα+ IELs or other T cell subsets in the lactating mammary gland (Fig. 4A). There were also no differences in previously described thymic progenitor subsets (PD-1+ or T-bet+) ^43^ in the lactating thymus when compared to nulliparous mice (Fig. S8A). This raises two possibilities: either thymic progenitors seed the mammary gland before 4 weeks of age and these few T cells expand into CD8αα+ IELs during pregnancy, or mammary CD8αα+ IELs are of extrathymic origin, potentially from other mucosal sites, and migrate to the mammary gland during late pregnancy. We found a modest increase in Ki67+ T cells during gestation, but not lactation, suggesting that the expansion of mammary T cells could be a combination of proliferation of mammary T cells and extrathymic input (Fig. S8B). We previously used Kaede photoconvertible mice to track the migration of immunocytes from the intestines to other body locations^44^. Among migratory intestinal cells in the spleen was a small population of IEL-like CD8αα+ TCRαβ+ cells, which led us to hypothesize that mammary CD8αα+ IELs could be of intestinal origin^44^. To test this hypothesis, intestinal sections (small intestine, excluding Peyer’s patches) of Kaede mice were photoconverted from green to red, at different times of gestation and early lactation, and mammary glands were analyzed 24hrs later (Fig. 4B). Importantly, due to the challenges associated with performing surgery in pregnant mice, only a small fraction of the intestine, accessible with minimal disturbance to surrounding tissues, was photoconverted. Kaede-red cells of intestinal origin had indeed migrated to the mammary gland, including all three CD8αα+, CD4+ and CD8αβ+ T cell types (Fig. 4C), and to the spleen as previously reported^44^ (Fig S8C). Although from small numbers, this observation indicates that some intestinal T cells migrated to the mammary gland within this one-day period. Thus, expansion of mammary T cells in late pregnancy and lactation is driven by thymic and intestinal inputs.

**Figure 4.**
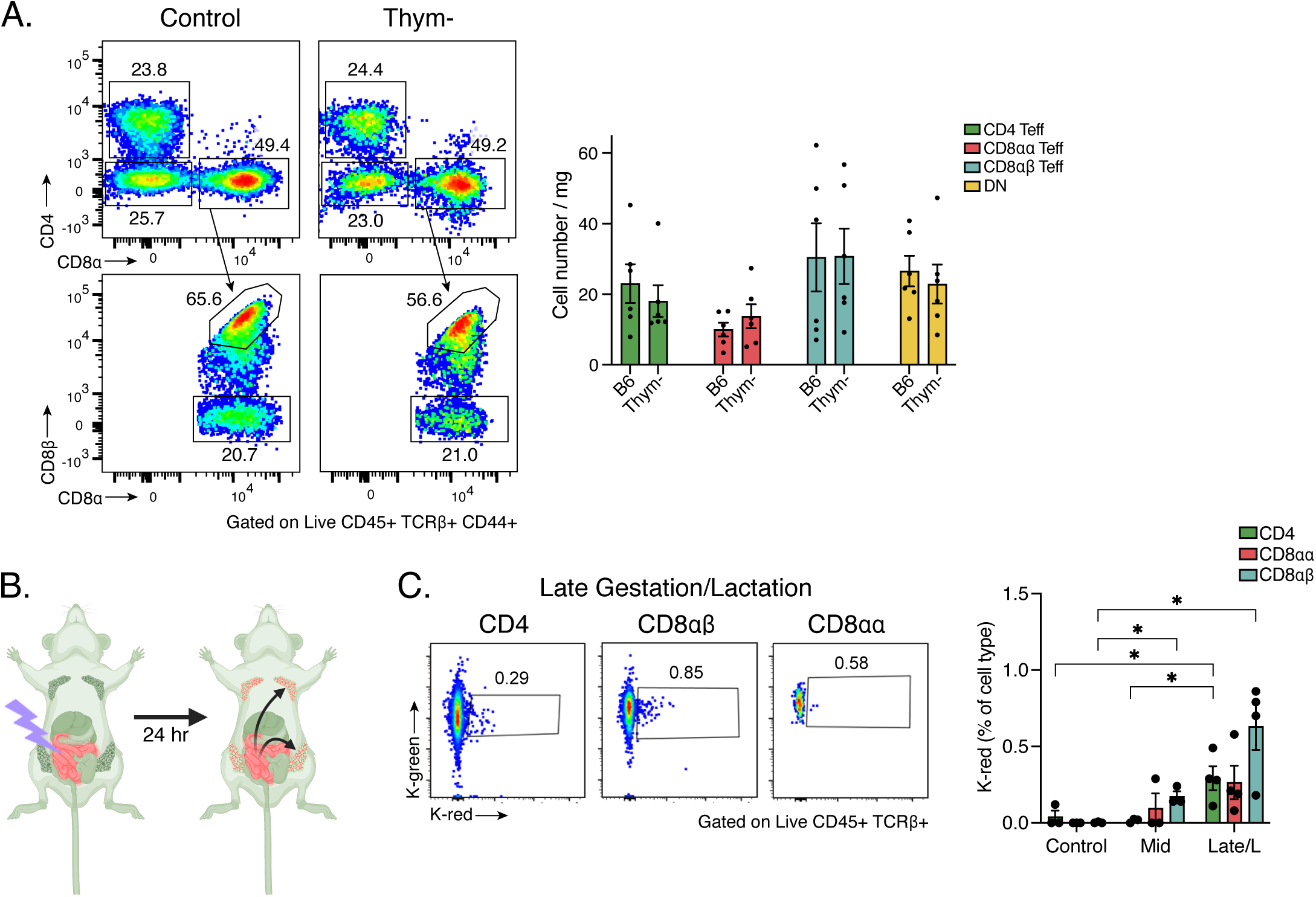
Intestinal T cells migrate to the mammary gland during late gestation and lactation. A) Representative flow cytometry plots and quantification of CD4 Teff, CD8⍺⍺+ Teff, CD8⍺β+ Teff and DN cell populations in control B6 (n=6) and thymectomized (n=6) lactating mice. Teff populations were determined as CD4 Teff: CD4+CD44+CD62L-. CD8αα Teff: CD8α+CD8β-CD44+CD62L-. CD8αβ Teff: CD8α+CD8β+CD44+CD62L-. DN: TCRβ+CD4-CD8α-. B) Experimental design for Kaede experiments: intestines of Kaede+ mice were photoconverted from green to red by illumination with UV light after laparotomy in mid-late pregnancy and early lactation, and migration of red cells to the mammary gland examined after 24 hours. C) Representative flow cytometry plots and quantification of Kaede red cells within T cell populations (gated on TCRβ+ followed by either CD4+ or CD8β+ or CD8⍺+ CD8β-) in the mammary gland of mice 24 hours post-photoconversion of the intestine. Controls are non-photoconverted mice (n=3), mid being mice photoconverted on gestation day 10 and analyzed on gestation day 11 (n=3) and late/L representing mice both photoconverted on gestation day 16 and analyzed on gestation day 17 and mice photoconverted on lactation day 1 and analyzed on lactation day 2 (n=4). * p<0.05 by two tailed student’s t-test (C). Data representative of ≥3 independent experiments, bars in plots indicate mean ± SEM.

### T cell clones are shared between intestinal IELs and mammary IELs

To further establish the relationship between intestinal and mammary IELs, we analyzed the αβTCR clonotypes expressed by T cells in the small and large intestine, mammary gland and spleen. First, we used single-cell TCRseq to compare αβTCR pairs displayed by IELs (small and large intestine) and mammary gland T cells in the same mice (four nulliparous and four lactating, 21,750 total cells). Overall, the data showed unremarkable V and J region usage, CDR3 length and N region diversity frequencies. Canonical TRAV11/TRAJ18 TCRs of iNKT cells were relatively abundant among lactating mammary T cells, mostly in CD4-CD8-DNs (2.2 and 8.1% of total cells, Fig. S9A). Rarefaction analysis revealed a notable degree of clonal amplification in different T cell-types from the lactating mammary gland compared to the nulliparous mammary gland; whereas amplification is seen in both the lactating and nulliparous small intestine (Fig. 5A, S9B), but with much mouse-to-mouse variation. We identified 13 TCR clonotypes shared between small intestine and mammary T cells in lactating mice, in contrast to 3 shared TCR clonotypes in nulliparous mice (Fig. 5B,C and Table S1). These shared clonotypes were defined by full nucleotide sequence identity, and were completely absent when comparing different mice, indicating that they stemmed from the same T cell clones present in both mammary gland and small intestine (and large intestine, for some). The clonotypes shared with the intestines accounted for 4.3% and 0.6% of mammary T cells in the two lactating mice, certainly an underestimate given incomplete sampling. As indicated in Fig. 5C and Table S1, clonotypes present in both small intestine and mammary gland belonged to several cell types, indicating that the exchange between tissues involves different T lineages. Some clonotypes shared between intestine and mammary gland were also observed in nulliparous females (2.2% and 0.9% of mammary T cells in the two mice profiled). Thus, exchanges of T cells between intestine and mammary gland pre-exist the onset of lactation.

**Figure 5.**
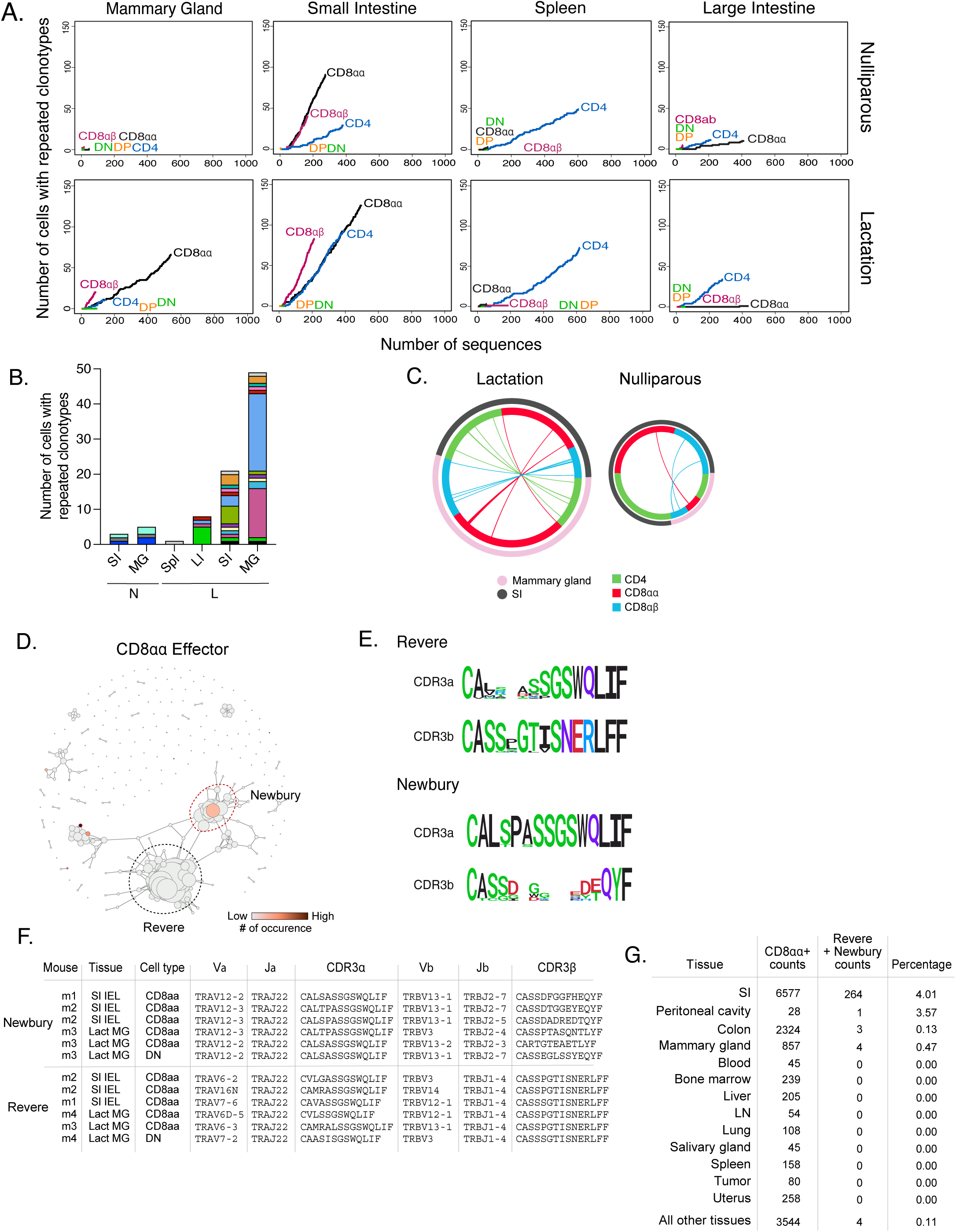
Mammary T cells share peculiar TCR repertoires with small intestinal T cells. A) Rarefaction analysis from TCR sequencing of T cell types between nulliparous and lactating mice in the mammary gland, small intestine, spleen and large intestine. B) Quantification of the number of cells with repeated clonotypes between organs in a nulliparous and lactating mouse. Each color representing a unique clonotype. C) Chord diagrams in lactating (L3) and nulliparous mice representing clonotype sharing in different T cell populations (inner ring) between the small intestine and mammary gland (outer ring). Each line represents a TCR clonotype. D) Distance matrix between αβTCR clonotypes in intestinal IELs. Red circle denotes “Newbury TCR” and black circle denotes “Revere TCR”. E) CDR3 sequence of Revere and Newbury TCRs. F) Table representing instances of Revere and Newbury TCRs in CD8⍺⍺+ T cells in mammary gland and small intestine across different mice. G) Counts of Revere and Newbury TCRs in CD8⍺⍺+ T cells across multiple tissues. Data representative of ≥3 independent experiments.

These clonotypic analyses also revealed the sharing of a particular intriguing group of cells. For a broader comparison of small intestine cells, we leveraged TCR sequence data of intestinal IELs generated in the ImmGenT program^45^, and used the TCRdist3 algorithm^46^ to compute a matrix of distances between αβTCR clonotypes. This revealed two prominent TCR families within the small intestinal IEL compartment (Fig. 5D), whose over-representation was striking because of the recruitment of highly related TCRs with little clonal amplification, as denoted by subtly different nucleotide sequences. These TCR families were found in many independent samples, and one indeed corresponded to the previously reported “Revere” family^47^ (we hereafter name the second family “Newbury” for consistency). Revere TCRs are mostly conserved in the CDR3b region, with exclusive usage of TRAJ22 and TRBJ1-4 (Fig. 5E, S10A), while the Newbury family is mostly conserved in the CDR3a region. Notably, these two families are almost exclusively represented in the intestinal CD8αα+ IELs, amounting to a few percent of T cells (Fig. S10B, C), and these highly identical TCRs of the CD8αα+ IELs are likely selected repeatedly by self-reactivity, in line with well-known selection of CD8αα differentiation by self-reactive transgenic TCRs^34, 35, 36^. Interestingly, Revere and Newbury TCRs were also observed in CD8αα+ IELs of the mammary gland (3 of each), with all the key sequence characteristics (Fig. 5F, S10D). Thus, peculiar TCR families of intestinal CD8αα+ IELs are found in IELs of the lactating mammary gland. Importantly, outside of the intestine and mammary gland, CD8αα+ IELs did not display Revere or Newbury family TCRs, as analyzed by the immgenT program (Fig. 5G, chisq.test p=6.10^-4^)). Together, CD8αα+ IELs in the lactating mammary gland displayed TCRs otherwise exclusive to intestinal IELs, as part of a broader exchange of T cell clones between the intestine and mammary gland.

### IEL-like cells are present in human mammary gland and human milk

We assessed whether mammary IELs were conserved across species by profiling T cells in human breast tissue and human milk. First, we used previously published scRNAseq datasets^48^ and found that human breast tissue from non-lactating women contains naïve (*SELL*) T cells as well as CD4+ and CD8αβ+ T cells that express tissue resident and cytotoxic markers expressed by mouse mammary IELs including *ITGAE*, *CD94*, *CD160*, *NKG2D,* and *GZMB* (Fig. 6A,B, S11A). We also observed a small population of cells that expressed genes associated with CD8αα+ IELs including *FCER1G* and *TYROBP.* The presence of CD4+ and CD8+ T cells in milk has been reported before^25^, but with no further characterization. To ask whether human milk contains CD8αα+ IEL-like cells in addition to the other subsets, we analyzed fresh milk samples from lactating women. Multiparameter flow cytometry revealed both CD4+ and CD8+ T cells in all samples. In addition, CD8αα+ IEL-like cells in human milk, including cells that expressed CD103, CD94, and NKG2D (Fig. 6C-E). Overall, we identified human counterparts of mouse mammary IELs in human breast and milk.

**Figure 6.**
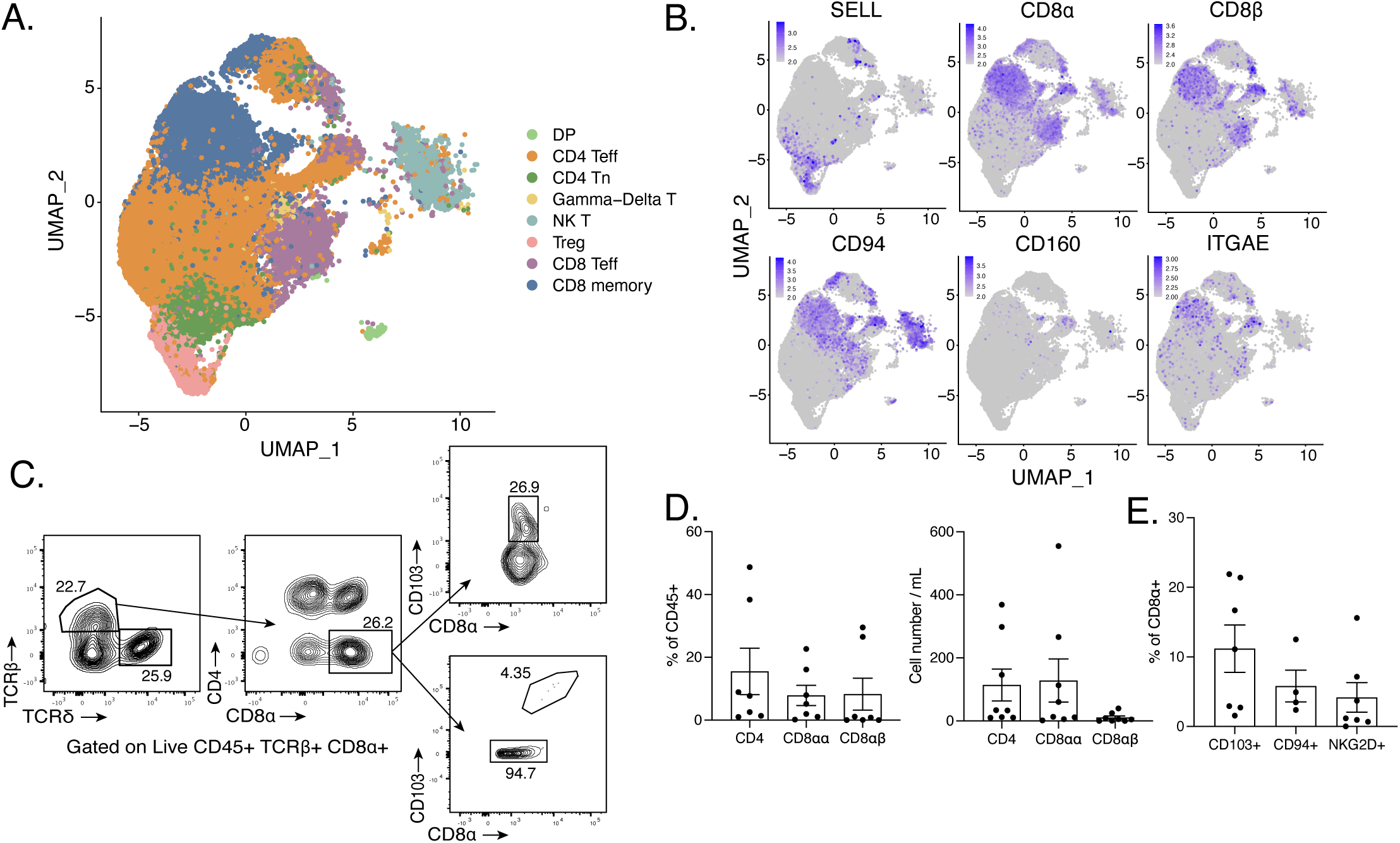
Mammary IEL-like cells are found in human breast and milk. A) UMAP projection of mammary immunocytes from human breast tissue (sourced from Kumar et al). B) Feature plots of selected genes projected on UMAP from (A). C) Representative flow cytometry gating of CD8α+ CD103+ T cells and CD8αα+ IEL-like cells in human milk samples. D) Quantification of CD4+, CD8αα+ and CD8αβ+ cells as percent of CD45+ cells (left) (n=7) and cell number normalized to volume (right) (n=8) in human milk samples. E) Proportion of human CD8⍺⍺ IEL-like cells that express markers CD103 (n=7), CD94 (n=4), and NKG2D (n=7) in human milk samples. Data represents ≥7 independent milk samples/experiments, bars in plots indicate mean ± SEM. Single cell sequencing from Kumar et al.

### Microbiota influence numbers and activation states of mammary IELs

The classical function of IELs is to maintain barrier immunity, which raised the question of microbe-dependence of IELs in the lactating mammary gland, considering the transition involves exposure to environmental microbes. Lactating mammary glands in microbe-deficient germ-free (GF) mice had morphologically different ducts compared to microbe-sufficient (specific pathogen free, SPF) control mice in hematoxylin-eosin stained sections (Fig. 7A). Although the number of mammary alveoli in SPF and GF mice were comparable, the average area (um^2^) of GF alveoli was larger, indicating that microbes may affect the developmental progression of the mammary gland during lactation (Fig. 7B, C). Microbes influenced total mammary immunocytes, as indicated by fewer total CD45+ cells in GF mice (Fig. 7D), but there were no differences in the proportion of basal or luminal epithelial cells between GF and SPF mice (Fig. S11B). There was also no significant difference in the average pup weight normalized to litter size (Fig. S11C), but whether microbes influence milk composition or production needs to be further investigated.

**Figure 7.**
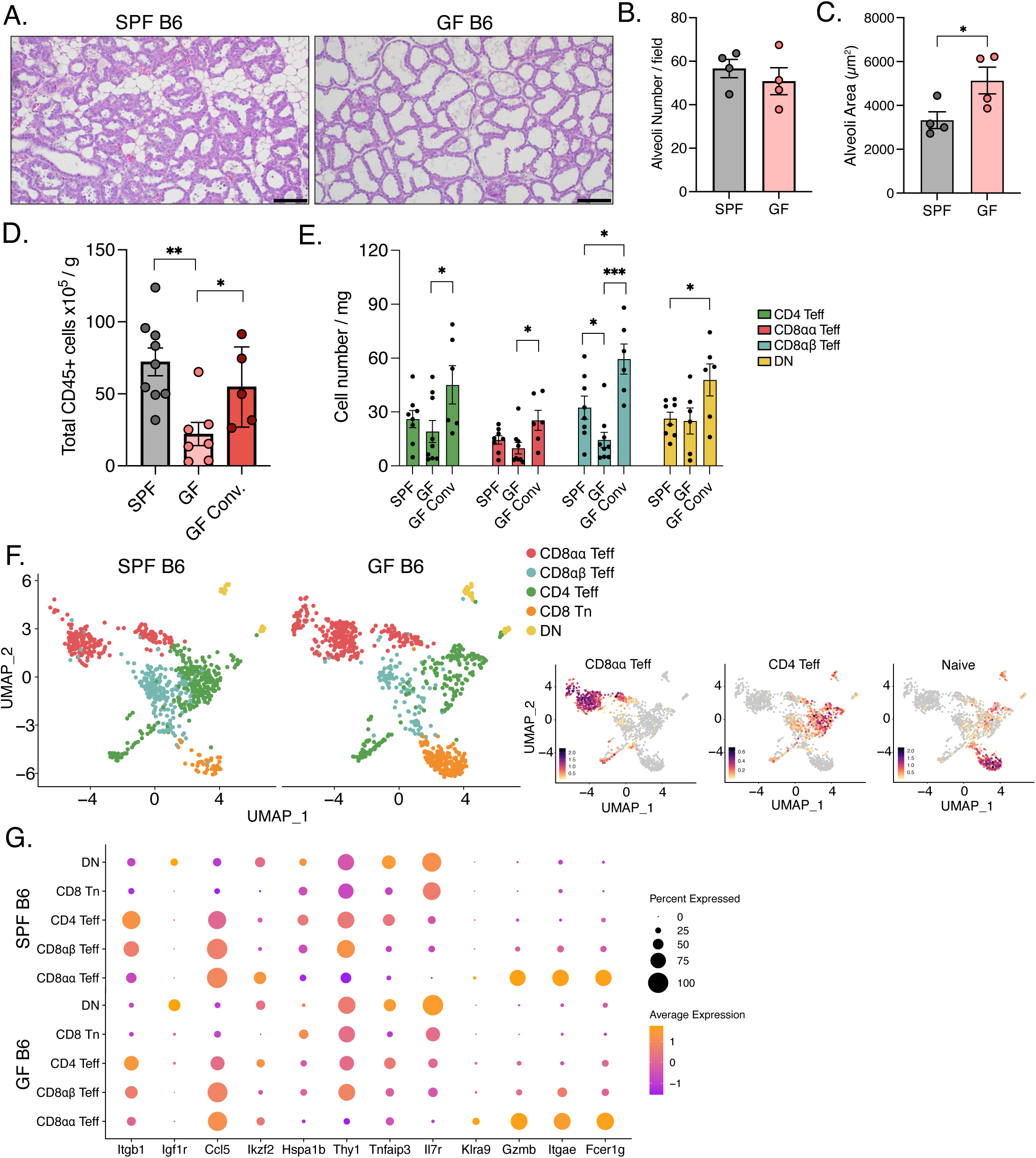
T cell expansion in the lactating mammary gland is partially dependent on microbes. A) Representative hematoxylin and eosin staining of lactating mammary gland of specific pathogen free (SPF) and germfree (GF) mice. Scale bar = 100µm. B, C) Quantification of the average number (B) and average area (µm^2^) (C) of alveoli per 20x image fields (five images per mouse of size 900µm x 500µm) of SPF and GF mammary glands (n=4). D) Quantification of the total number of CD45+ cells normalized to mammary gland weight in SPF (n=9), GF (n=7) and GF conventionalized mice (n=5). E) Quantification of total cell numbers normalized to mammary gland weight, of specified T cell populations in SPF (n=8), GF (n=9, n=6 for DN) and GF conventionalized (n=6) mice by flow cytometry. Teff populations were determined as CD4 Teff: CD4+CD44+CD62L-. CD8αα Teff: CD8α+CD8β-CD44+CD62L-. CD8αβ Teff: CD8α+CD8β+CD44+CD62L-. DN: TCRβ+CD4-CD8α-. F) UMAP projection of mammary T cells from lactating SPF and GF mice with feature plots of naïve and effector T cell gene signatures. G) Dot plot of differentially expressed genes in mammary T cell populations between lactating SPF and GF mice from (F). *p<0.05, **p<0.01 ***p<0.001 by student’s t-test. Data representative of ≥3 independent experiments, bars in plots indicate mean ± SEM.

GF mice showed decreased numbers of CD4+ Teff, CD8αα+ Teff, and CD8αβ+ IELs compared to SPF mice (Fig. 7E). However, the proportions of T cell types were not different between the groups suggesting that the decrease in IEL numbers was due to the total drop in CD45+ cells (Fig. S11D). The decrease in mammary IELs could also stem from decreased intestinal IELs, as GF mice display a significant decrease in CD4+ IELs and CD8αβ+ IELs, and a modest decrease in intestinal CD8αα+ IELs^49^. The defect in mammary IELs may arise from migrating intestinal IELs that lack the same functionality, reflecting the role of intestinal microbes in priming mammary IELs. To test the role of intestinal microbes, we conventionalized GF mice by transferring fecal microbes from control SPF mice into GF mice at 6 weeks of age. Conventionalized GF mice displayed increased CD45+ cells during lactation, including restoration of the mammary IEL populations (Fig. 7D,E). To further analyze micro-dependent phenotypic changes, we performed multiplexed scRNAseq on T cells from lactating mammary glands from GF or SPF mice (Fig. 7F, 13,021 cells, 8 mice, 2 independent runs). All mammary IEL populations were equally represented in the two groups consistent with quantitation by flow cytometry (Fig. 7F). Differential gene expression between SPF and GF IELs showed few transcriptional differences, including significant changes in *Itgb1, Igf1r, Ikzf2, Tnfaip3, Thy1*, and *Klra9* (Fig. 7G). While several of these genes are immunomodulatory, whether these changes affect the function of mammary IELs in GF mice needs to be examined. Thus, our data points to the importance of commensal microbes in modulating numbers of mammary gland IELs during lactation.

## DISCUSSION

We report a dynamic atlas of T cells whose changes accompany the adaptation of the mammary gland to a mucosal state in lactation, with an increase in activated CD4+, CD8αβ+ and CD8αα+ T cells that are intraepithelial in location. Mammary CD8αα+ IELs shared TCR clonotypes with intestinal CD8αα+ TCRαβ+ IELs, suggesting T cell migration from the intestine to the mammary gland. Mammary IELs may be conserved across species since human breast tissue and milk contained T cells with a similar signature. Finally, we found that T cell numbers and activation in lactation were partly influenced by microbes, suggesting that the observed T cell changes could be geared towards promoting barrier immunity during lactation.

Intestinal IEL subsets maintain and protect the intestinal barrier^31^, functions that mammary IELs could be involved in to promote lactogenesis. The TCR specificity of CD8αα+ IELs is directed towards self-antigens^34, 35, 36^, and in line with that, we found two families of TCRs that are repeatedly generated and selected across mice, and were found in the small intestine and lactating mammary gland. CD8αα homodimers have been shown to function as TCR corepressors by binding to TL (thymus leukemia) antigen on epithelial cells, to negatively regulate T cell activation by decreasing antigen sensitivity, in contrast to CD8αβ heterodimers that enhance TCR function^50^. Our results suggest that mammary CD8αα+ IELs are poised to respond to the rapid epithelial cell proliferation, or to the plethora of self-antigens that are present in the lactating mammary gland, but whether their function is tolerogenic or cytotoxic needs to be determined. During pregnancy and lactation, the expression of gut-homing markers Ccl25, Ccl28, and MAdCAM1, increase in the mammary tissue which could lead to T cell migration to the mammary gland^22, 51, 52^. However, whether the expression of these homing markers change in the intestine during pregnancy and lactation which mediate T cell egress, and whether hormones influence mammary homing markers needs to be further explored.

Apart from their physiological role in lactogenesis, CD8αα+ IELs may influence post-lactation oncogenesis. Similar populations of NK-like unconventional T cells such as αβILTCKs and NK-like ψο T cells, have been shown to be important in suppressing mammary oncogenesis^26, 32^. Mammary T cells are present in human milk where function, if any, is unclear. One possibility is that their presence is passive, and linked to epithelial cell sloughing in lactogenesis, which increases epithelium-associated T cells in milk. Maternal T cells have been previously suggested to colonize neonatal intestines^53^, raising the other possibility that milk IELs could migrate into and colonize the neonatal intestine to promote barrier protection.

Pregnancy induces an immunosuppressive state to maintain maternal-fetal tolerance and support successful pregnancy outcomes^54^. However, in the mammary gland, we observe an increase in effector T cell populations in late pregnancy and early lactation presumably to prepare for increased microbial exposure or increased epithelial cell proliferation in lactation. Consistent with the idea of increased microbial exposure, activated T cells decrease in the GF mammary gland, and the drop in mammary Tregs in lactating SPF mice is not observed in GF mice (Fig. S11E). Interestingly, liMacs were also reduced in lactating GF mammary glands^17^, suggesting that microbes or microbe-derived signals can influence multiple immunocyte-types involved in mammary remodeling.

In summary, we have characterized novel T cell changes during lactogenesis and provide evidence for T cell migration potentially along the entero-mammary axis. Our results set the stage for deepening our understanding of T cell function in lactogenesis, which could provide new strategies to improve maternal defense and tolerance during and after lactation.

## METHODS

### Mice

C57BL/6 (B6) mice were purchased from Jackson Labs and maintained in specific pathogen free (SPF) conditions at Harvard Medical School and Salk Institute for Biological Sciences. Nulliparous mice were littermate controls of mice profiled at pregnancy, lactation, or involution. For timed pregnancies, female B6 mice were set up at 6-8 wks of age with male B6 mice, female mice with plugs were separated and housed individually for the duration of pregnancy, and mammary glands were profiled at day 12 (G12), day 17 (G17) of pregnancy, lactation day 3-5 (L3), and involution day 1 (I), one day post-weaning of pups at day 21. All experiments were performed following guidelines listed in animal protocols (IS00001257, Harvard Medical School) and (23-00007, Salk Institute for Biological Studies) approved by the Institutional Animal Care and Use Committee.

Germ free (GF) B6 mice were purchased at timepoints listed above from the University of California San Diego. GF mice were conventionalized by oral gavage of fecal microbiota from SPF B6 mice, one week prior to mating and maintained in SPF conditions. Kaede reporter mice were obtained from O. Kanagawa (RIKEN, Wako, Japan) and maintained on the B6 background^55, 56^.

### Preparation of lymphocytes and flow cytometry

#### Mammary gland

Inguinal lymph nodes were removed and mammary glands 3, 4 and 5 were collected, minced and dissociated in collagenase solution (3mg/mL collagenase type II (Sigma C6885) and 2% FBS in DMEM) in a 37°C shaking water bath for 20 min with manual shaking every 5 min, followed by red blood cell lysis. Single cell suspensions were filtered and washed with 2% DMEM solution.

#### Thymus and LN

Lymphocytes from thymus and inguinal lymph nodes were obtained by mechanical disruption, filtered and washed with 10% RPMI solution.

#### Intestines

Small and large intestinal tissues were measured, cleaned, and treated with RPMI containing 1mM DTT, 20mM EDTA and 2% FBS at 37°C for 15 min to isolate the epithelial and IEL fractions. For the lamina propria (LP) fraction, the remaining tissue was dissociated in collagenase solution (1 mg/mL collagenase VIII (Sigma C2139), 50μg/ml DNase (Sigma C6885) in 1%FBS in RPMI) with constant stirring at 37°C for 30min. Single cell suspensions for the IEL and LP fractions were filtered and washed with 10% RPMI solution.

#### Spleen

Tissue was mechanically disrupted, followed by red blood cell lysis. Single cell suspensions were filtered and washed with 10% RPMI solution.

#### Staining

Single cell suspensions of cells resulting from tissue dissociations were stained with different panels of antibodies with surface markers for CD45, CD4, CD8α, CD8β, TCRβ, TCRδ, NK1.1, Ly49, CD103, Thy1, PD-1, CD122, CD5, CD69, CD44, CD62L, CD38, CD244, and CD160 Zombie UV Fixable Viability and intracellular markers for T-bet, Ki67 and Foxp3. For intracellular staining, cells were stained for surface markers and fixed in eBioscience Fix/Perm buffer overnight, followed by permeabilization in eBioscience permeabilization buffer at room temperature for 45 min in the presence of antibodies. Cells were acquired with a BD LSRII or BD FACSymphony A3 and analysis was performed with FlowJo 10 software.

### Photoconversion Procedure

Kaede transgenic mice were anesthetized, abdomen was surgically opened and a portion (approximately one third in non-pregnant mice, smaller portion in pregnant mice) of the small intestine was exposed. The mouse, except for the small intestine, was covered in aluminum foil and the small intestine was exposed to a handheld 405 nm blue purple laser for 30 second light pulses (which converts Kaede green cells to Kaede red). After photoconversion the mouse was surgically closed and sacrificed 24 hours later for flow cytometry analysis of Kaede green vs Kaede red cells.

### Single cell RNA and TCR sequencing

#### Mammary immunocytes

Live CD45+ cells were sorted from the mammary gland of nulliparous (n=6), gestation day 17 (n=4), lactation day 3-5 (n=6), and involution (n=4) mice using a BD FACSAria after hashtagging with Biolegend TotalSeq-A reagents, and samples were pooled for encapsulation (10X Chromium). Libraries were prepared using Chromium Single cell 3’ reagents kit v2 and sequenced on NovaSeq 6000.

#### Multi-organ combined scRNAseq and TCRseq

Live T cells (DAPI-CD3+CD44+TCRβ+) were sorted from the mammary gland, small intestines, large intestines, spleen and thymus from nulliparous (n=4) and lactation day 4 (n=4) mice. The cells were hashtagged with Biolegend TotalSeq-C reagents and pooled for encapsulation (10X Chromium). Libraries were prepared using Chromium Single cell 5’ reagents kit v3 and sequenced on NovaSeq 6000. TCR and hashtag libraries were processed as described^29^.

#### Germ-free vs SPF

Live T cells (DAPI-CD3+CD44+TCRβ+) were sorted from GF (n=4) and SPF (n=4) mammary glands on lactation day 4. Samples were pooled for encapsulation (10X Chromium), libraries were prepared using Chromium Single cell 3’ reagents kit v3 and sequenced on NovaSeq 6000.

#### Epithelial-IEL interactions

Live EpCAM+ CD45-, CD45+ EpCAM-cells, and TCRβ+ cells were sorted, pooled for encapsulation (10X Chromium), libraries prepared using Chromium Single cell 3’ reagents kit v3 and sequenced on NovaSeq 6000.

Single-cell RNAseq data was analyzed using the Seurat pipeline, which allowed for data normalization, clustering, and identification of differentially expressed genes across groups.

### Cell Interaction Predictions

CellChat v2 was used to infer and visualize intercellular communication networks in the mammary gland^40, 57^. CellChat v2 is an R package that is able to predict and analyze intercellular communication pathways from single-cell data. The analysis was done as described in the CellChat v2 published protocol. Briefly, EpCAM+ and TCRβ+ cells were isolated from the scRNA-seq data and used to predict intercellular communication pathways. To perform comparison analysis, we isolated differentially expressed ligands and receptors between nulliparous and lactating mice (p < 0.05), which were used to predict communication pathways that could be different between these groups. For visualization purposes, the networks between EpCAM+ and TCRβ+ cells were isolated and visualized using chord diagrams. Summary of signaling pathways were generated using CellChat v2 and visualized using chord diagrams.

### Histology, Imaging and Microscopy

Mammary gland 4 was harvested from nulliparous, gestation (G17), lactation (L3-5), and involution (I) stages and fixed in 4% Paraformaldehyde (PFA) solution in PBS overnight at 4°C shaking. They were washed with PBS and stored in 70% ethanol prior to being embedded in paraffin. Immunofluorescence staining was performed as previously described^58^. All primary antibodies were diluted in Renaissance Background reducing diluent (Biocare, PD905L). All opals were diluted 1:500 in 1X Plus Manual Amplification Diluent (Akoya Biosciences, FP1498).

Microscopy methods are reported following the guidance of (Montero-llopis et al, 2021)^59^ for best reproducible practices. Images in Figures 2E and S4D were acquired using an Olympus VS200 Slide Scanner widefield microscope equipped with a NOCEM X-cite light source (405-780nm) and the fluorescent camera Hamamatsu Orca fusion BTsCMOS (2304×2304 pixels 6.5 um). Images were acquired using a UPlan X Apo 20x/0.8 air objective. Signal from DAPI, FITC, TRITC, CY5 and CY7 was collected by illuminating the sample using the FF409/493/573/652-Di02 or FF757-Di01 multiband dichroics and the following excitation (FF01-378/52, FF01-474/27, FF01-554/23, FF01-635/18, FF01-735/28) and emission (FF01-432/36, FF01-515/30, FF01-595/31, FF01-698/70, FF02-809/81) filters respectively. Images were acquired using the OlyVIA software from Olympus and processed by Qpath and FIJI to crop representative areas and threshold background signal.

For H&E staining, sections were deparaffinized, stained with hematoxylin and eosin, dehydrated and mounted with coverslips and imaged on an Olympus upright INSERT SCOPE brightfield microscope at 10x and 20x magnification.

#### Milk Alevoli Quantification

Milk duct area was measured using Fiji to measure each duct in 4 different 20x magnification images per mouse and the average of all ducts was calculated.

### Human Milk

5 mL of human milk were received from UC San Diego HMB Biorepository and diluted 1:1 with PBS. Milk samples were centrifuged to remove lipid and whey layers. Remaining cells were stained using viability dye, CD4, CD94, CD8β, CD3, TCRδ, CD44, CD45, CD8α, CD103, and NKG2D. Cells were analyzed using BD LSRFortessa or BD FACSymphony A3 and analysis was performed with FlowJo 10 software. The protocol for analysis of human milk samples was approved by the UCSD Institutional Review Board Administration (IRB number: 808920) and patients provided written informed consent prior to enrollment.

### Quantification and Statistical Analysis

Data are presented as mean ± SEM. Unless stated otherwise, significance was assessed by Student’s t-test or one-way Anova in GraphPad Prism 8.0.

## Data availability

Single cell sequencing data are available in NCBI with accession numbers GSE290256 and GSE288901. Mammary gland T cell data are available in a user-friendly format at https://cbdm.connect.hms.harvard.edu/ImmgenT/PublicRosetta/

## ACKNOWLEDGEMENTS

We would like to thank Gabriel Ascui-Gac and Dr. Mitchell Kronenburg for reagents and advise on MAIT cell analysis, and Dr. Queralt Vallmajo Martin and Dr. Danish Umar for helpful discussions. This work was supported by NIH grants RO1-AI150686 to CB, R24-072073 to the ImmGen Consortium, NIH-NCI CCSG: P30 CA01495, and Shared Instrumentation Grants S10-OD023689 to the Salk Institute core facilities (FCCF, NGS).

## ImmgenT consortium members

David Zemmour, Christophe Benoist, Dania Mallah, Ian Magill, Liang Yang, Ananda Goldrath, Joonsoo Kang, Mitch Kronenberg, Nika Abdollahi, Myriam Croze, Serge Candéias, Sofia Kossida, Véronique Giuducelli, Sam Behar, Remy Bosselut, Laurent Brossay, Ken Cadwell, Alexander Chervonsky, Laurent Gapin, Jun Huh, Iliyan Iliev, Bana Jabri, Steve Jameson, Marc Jenkins, Susan Kaech, Dan Kaplan, Vijay Kuchroo, Stefan Muljo, Michel Nussenzweig, Marion Pepper, David Sinclair, Michael Starnbach, Paul Thomas, Ulrich von Andrian, Derek Bangs, Olga Barreiro del Rio, Gavyn Bee, Katharine Block, Sam Borys, Evelyn Chang, Josh Choi, Enxhi Ferraj, Giovanni Galletti, Lizzie Garcia-Rivera, Anna-Maria Globig, Xutong Gong, Takato Kusakabe, Rocky Lai, Andrea Lebron-Figueroa, Woan-Yu Lin, Mariya London, Erin Lucas, Ankit Malik, Julia Merkenschlager, Nathan Morris, Kevin Osum, Jinseok Park, Val Piekarsa, Sara Quon, Shanelle Reilly, Stefan Schattgen, Tomoyo Shinkawa, Nalat Siwapornchai, Claire Thefane, Melanie Vacchio, Jordan Voisine, Eric Weiss, Alia Welsh, Shanshan Zhang.

