## Supplemental Figures 1-11 for "Expansion of mammary intraepithelial lymphocytes and intestinal inputs shape T cell dynamics in lactogenesis"

### Figure S1

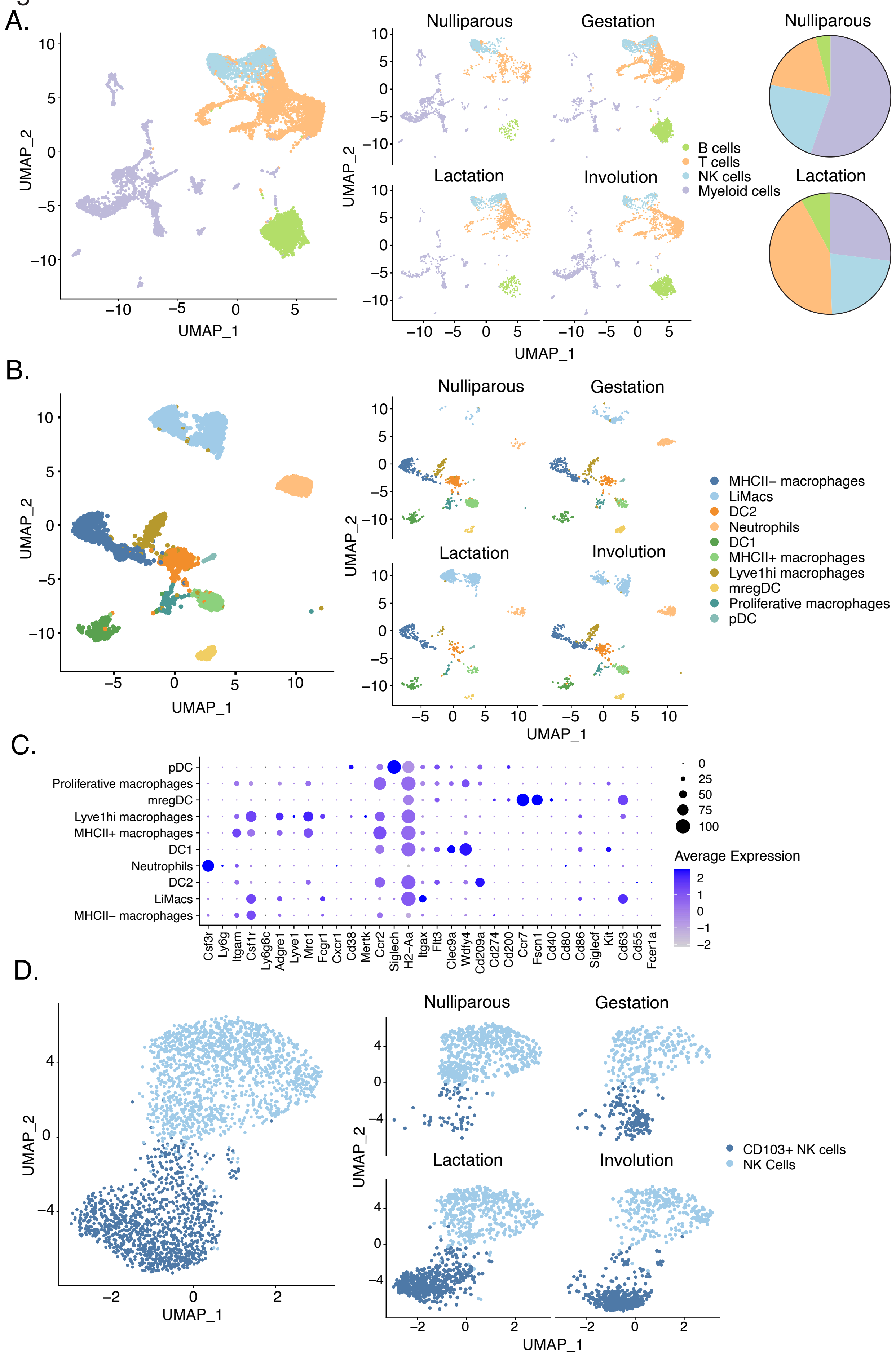

A.
UMAP_1
UMAP_2
Nulliparous
Gestation
Lactation
Involution
B cells
T cells
NK cells
Myeloid cells
B.
MHCII– macrophages
LiMacs
DC2
Neutrophils
DC1
MHCII+ macrophages
Lyve1hi macrophages
mregDC
Proliferative macrophages
pDC
C.
Average Expression
Csf3r
Ly6g
Itgam
Csf1r
Ly6g6c
Adgre1
Lyve1
Mrc1
Fcgr1
Cxcr1
Cd38
Mertk
Ccr2
Siglech
H2–Aa
Itgax
Flt3
Clec9a
Wdfy4
Cd209a
Cd274
Cd200
Ccr7
Fscn1
Cd40
Cd80
Cd86
Siglecf
Kit
Cd63
Cd55
Fcer1a
D.
CD103+ NK cells
NK Cells

**Supplementary Figure 1. Myeloid populations in the mammary glands during gestation and lactation.**

A) UMAP projection of all immunocytes in the mammary gland (left) split by stages nulliparous, gestation (G17), lactation (L3), and involution (right). Pie charts summarizing the proportions of cell populations in each stage (far right).

B) UMAP projection of all myeloid cells in the mammary gland (left) split by stages nulliparous, gestation (G17), lactation (L3), and involution (right).

C) Dot plot of myeloid cell population markers. Dot size represents the percentage of cells expressing the selected gene and color indicates expression level.

D) UMAP projection of NK cell populations in the mammary gland (left) split by stages nulliparous, gestation (G17), lactation (L3), and involution (right).

Data representative of 3 independent experiments.

### Figure S2

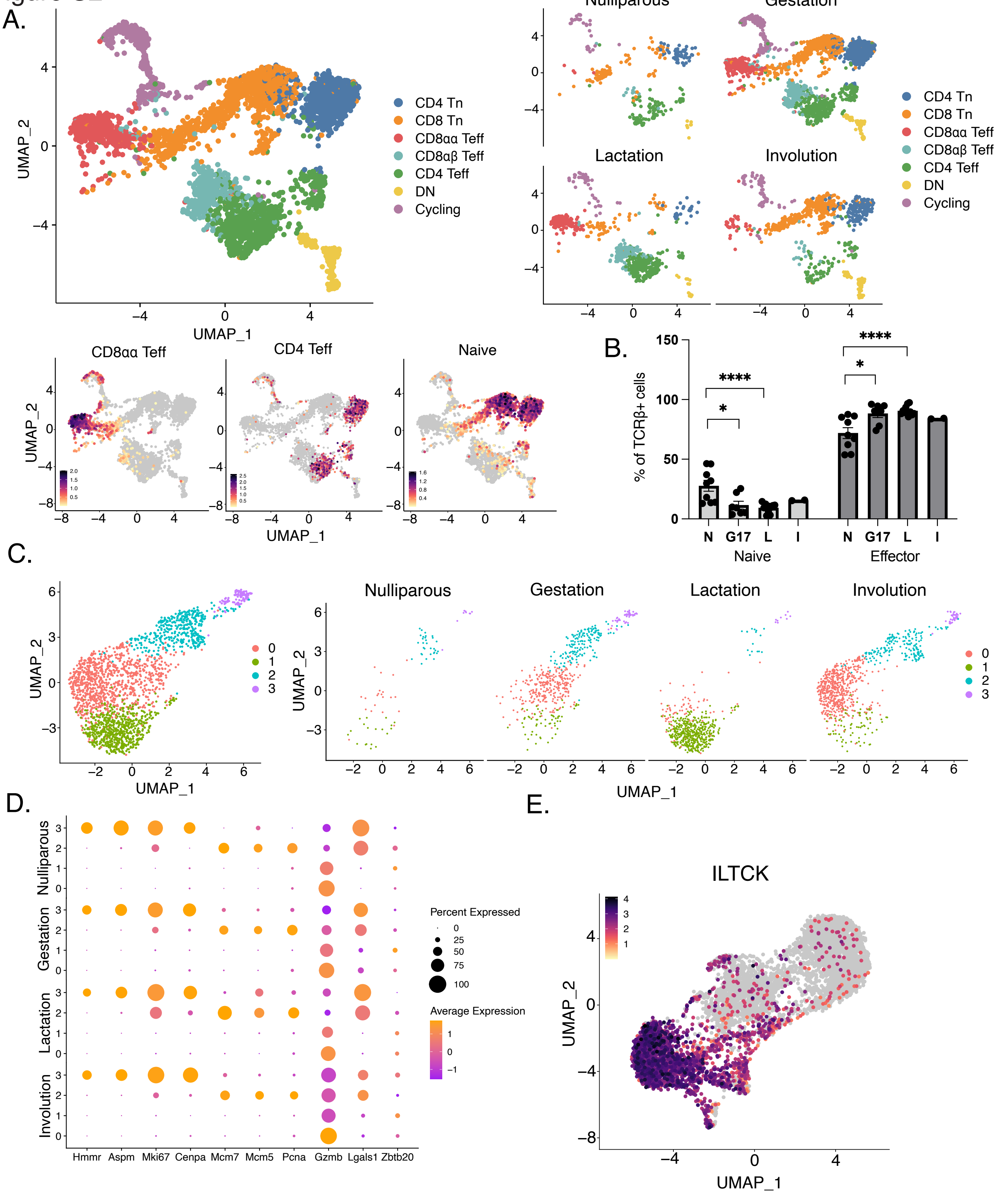

A.
UMAP_1
UMAP_2
CD4 Tn
CD8 Tn
CD8αα Teff
CD8αβ Teff
CD4 Teff
DN
Cycling
Nulliparous
Gestation
Lactation
Involution
CD8αα Teff
CD4 Teff
Naive
B.
% of TCRβ+ cells
N
G17
L
I
Naive
Effector
C.
0
1
2
3
D.
Hmmr
Aspm
Mki67
Cenpa
Mcm7
Mcm5
Pcna
Gzmb
Lgals1
Zbtb20
Percent Expressed
Average Expression
E.
ILTCK

**Supplementary Figure 2**. **T cell changes in the lactating mammary gland are reproducible across mice and experiments.**

A) UMAP projection of mammary T cells with feature plots of specified T cell gene signatures (left). Split by stages: nulliparous, gestation (G17), lactation (L3), and involution (right).

B) Quantification of the proportion of CD44+ (effector) and CD44- (naive) T cells across stages. N=nulliparous (n=9), G17=gestation day 17 (n=7), L=lactation days 3-5 (n=11) and I=involution, 1 day post weaning (n=2).

C) UMAP projection of CD8αα+ cell populations (left) split by stages nulliparous, gestation (G17), lactation (L3), and involution (right).

D) Dot plot of selected highly upregulated genes for CD8αα + populations across stages identified in (C). Dot size represents the percentage of cells expressing the selected gene and color indicates expression level.

E) UMAP projection of the ILTCK gene signature (Chou et al. 2022) on the summary T cell UMAP from Figure 1C.

*$p<0.05$, ****$p<0.0001$ by student's t-test. Data representative of ≥3 independent experiments, bars in plots indicate mean ± SEM.

Figure S3

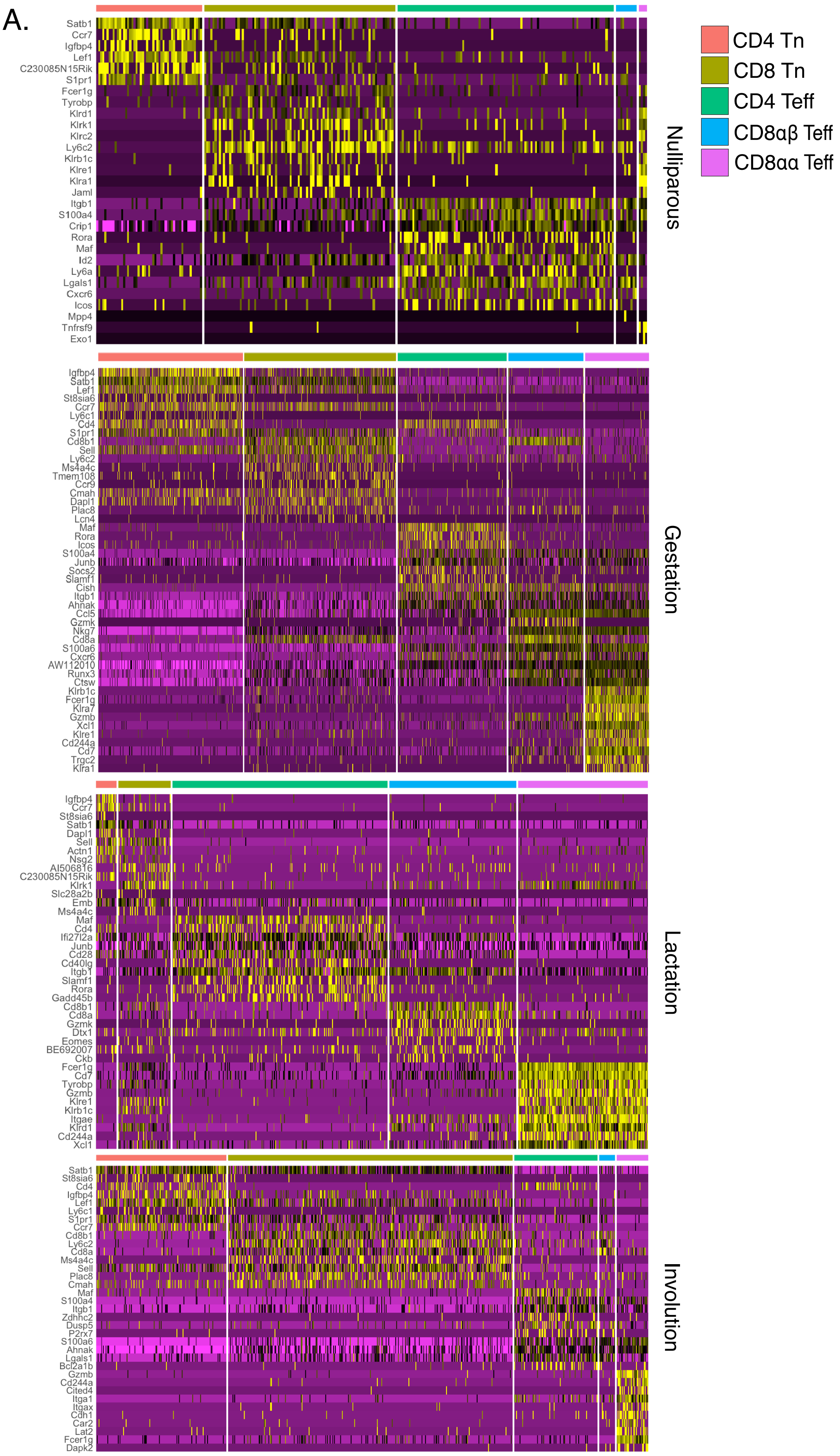
A.
CD4 Tn
CD8 Tn
CD4 Teff
CD8αβ Teff
CD8αα Teff
Nulliparous
Satb1
Ccr7
Igfbp4
Lef1
C230085N15Rik
S1pr1
Fcer1g
Tyrobp
Klrd1
Klrk1
Klrc2
Ly6c2
Klrb1c
Klre1
Klra1
Jaml
Itgb1
S100a4
Crip1
Rora
Maf
Id2
Ly6a
Lgals1
Cxcr6
Icos
Mpp4
Tnfrsf9
Exo1
Gestation
Igfbp4
Satb1
Lef1
St8sia6
Ccr7
Ly6c1
Cd4
S1pr1
Cd8b1
Sell
Ly6c2
Ms4a4c
Tmem108
Ccr9
Cmah
Dapl1
Plac8
Lcn4
Maf
Rora
Icos
S100a4
Junb
Socs2
Slamf1
Cish
Itgb1
Ahnak
Ccl5
Gzmk
Nkg7
Cd8a
S100a6
Cxcr6
AW112010
Runx3
Ctsw
Klrb1c
Fcer1g
Klra7
Gzmb
Xcl1
Klre1
Cd244a
Cd7
Trgc2
Klra1
Lactation
Igfbp4
Ccr7
St8sia6
Satb1
Dapl1
Sell
Actn1
Nsg2
AI506816
C230085N15Rik
Klrk1
Slc28a2b
Emb
Ms4a4c
Maf
Cd4
Ifi27l2a
Junb
Cd28
Cd40lg
Itgb1
Slamf1
Rora
Gadd45b
Cd8b1
Cd8a
Gzmk
Dtx1
Eomes
BE692007
Ckb
Fcer1g
Cd7
Tyrobp
Gzmb
Klre1
Klrb1c
Itgae
Klrd1
Cd244a
Xcl1
Involution
Satb1
St8sia6
Cd4
Igfbp4
Lef1
Ly6c1
S1pr1
Ccr7
Cd8b1
Ly6c2
Cd8a
Ms4a4c
Sell
Plac8
Cmah
Maf
S100a4
Itgb1
Zdhhc2
Dusp5
P2rx7
S100a6
Ahnak
Lgals1
Bcl2a1b
Gzmb
Cd244a
Cited4
Itga1
Itgax
Cdh1
Car2
Lat2
Fcer1g
Dapk2

**Supplementary Figure 3. Differential gene expression in T cell populations across stages.**

A) Heatmaps of top differentially expressed genes in specified T cell populations across nulliparous, gestation, lactation and involution stages.

Data is representative of 3 independent experiments.

Figure S4

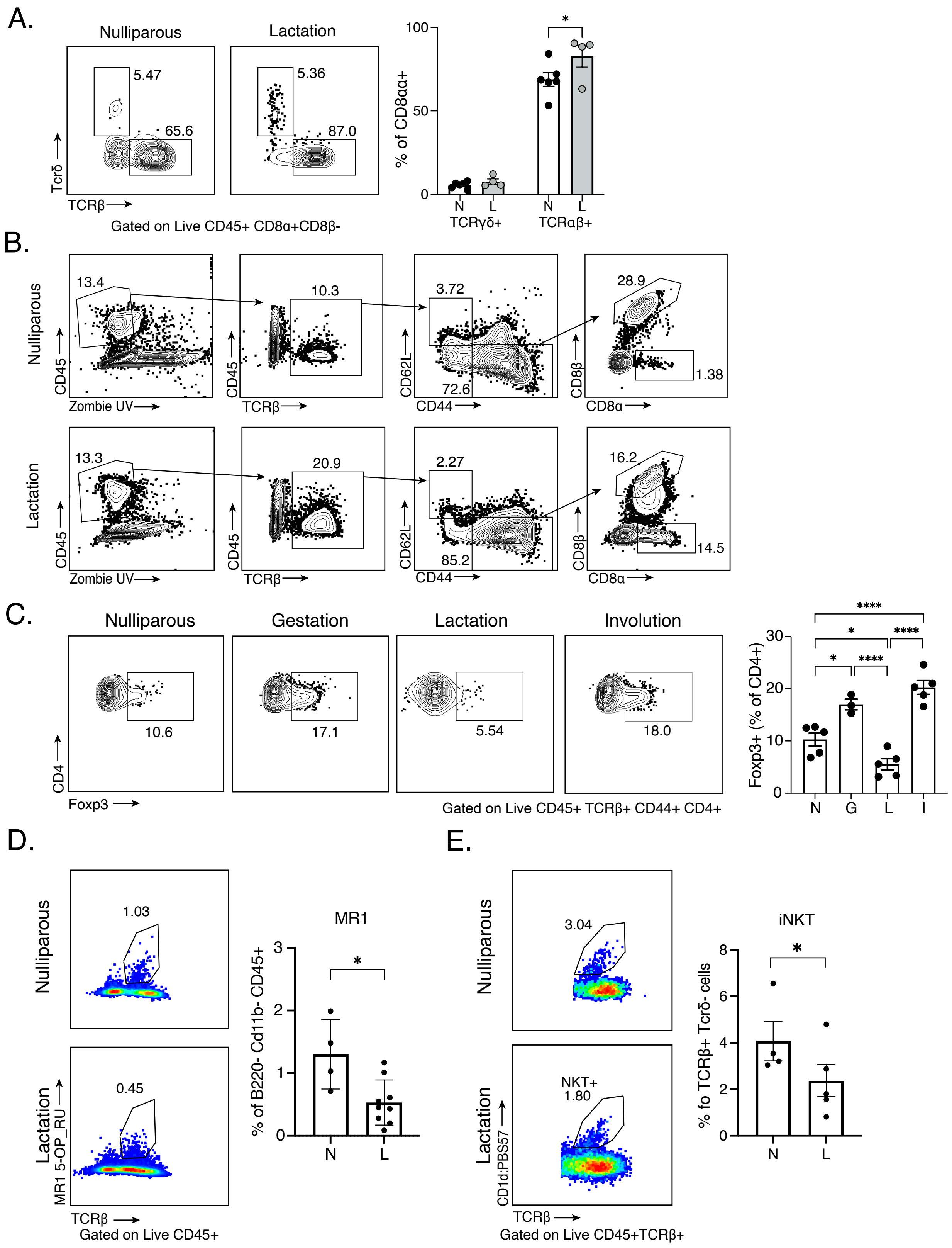
A.
Nulliparous
Lactation
5.47
65.6
5.36
87.0
Tcrδ
TCRβ
Gated on Live CD45+ CD8α+CD8β-
% of CD8αα+
100
50
0
*
N
L
N
L
TCRγδ+
TCRαβ+
B.
Nulliparous
Lactation
13.4
10.3
3.72
72.6
28.9
1.38
13.3
20.9
2.27
85.2
16.2
14.5
CD45
Zombie UV
TCRβ
CD62L
CD44
CD8β
CD8α
C.
Nulliparous
Gestation
Lactation
Involution
10.6
17.1
5.54
18.0
CD4
Foxp3
Gated on Live CD45+ TCRβ+ CD44+ CD4+
Foxp3+ (% of CD4+)
30
20
10
0
****
*
****
*
****
N
G
L
I
D.
Nulliparous
Lactation
1.03
0.45
MR1 5-OP_RU
TCRβ
Gated on Live CD45+
MR1
% of B220- Cd11b- CD45+
3
2
1
0
*
N
L
E.
Nulliparous
Lactation
3.04
NKT+
1.80
CD1d:PBS57
TCRβ
Gated on Live CD45+TCRβ+
iNKT
% fo TCRβ+ Tcrδ- cells
8
6
4
2
0
*
N
L

**Supplementary Figure 4. Unconventional T cell subsets are a minor population in the mammary gland.**

A) Representative flow cytometry plots and quantification indicating the proportion of mammary CD8αα+ cells that are TCRδ+ and TCRβ+ in N=nulliparous (n=6) and L=lactation days 3-5 (n=4) mice.

B) Representative gating for flow cytometry analysis with final gating on the CD8αα+ and CD8αβ+ populations in nulliparous and lactation.

C) Representative flow cytometry plots and proportion of mammary Foxp3+ T regulatory cells in N=nulliparous (n=5), G=gestation day 17 (n=3), L=lactation days 3-5 (n=5) and I=involution, 1 day post weaning (n=5).

D) Representative flow cytometry plots and proportion of mammary MR1 5-OP_RU+ T cells of CD45+ cells in N=nulliparous (n=4) and L=lactation days 3-5 (n=9) mice.

E) Representative flow cytometry plots and proportion of mammary iNKT+ cells of all CD45+ TCRβ+ cells in N=nulliparous (n=4) and L=lactation days 3-5 (n=5) mice.

*$p<0.05$, ****$p<0.0001$ by student's t-test. Data representative of ≥3 independent experiments, bars in plots indicate mean ± SEM.

Figure S5

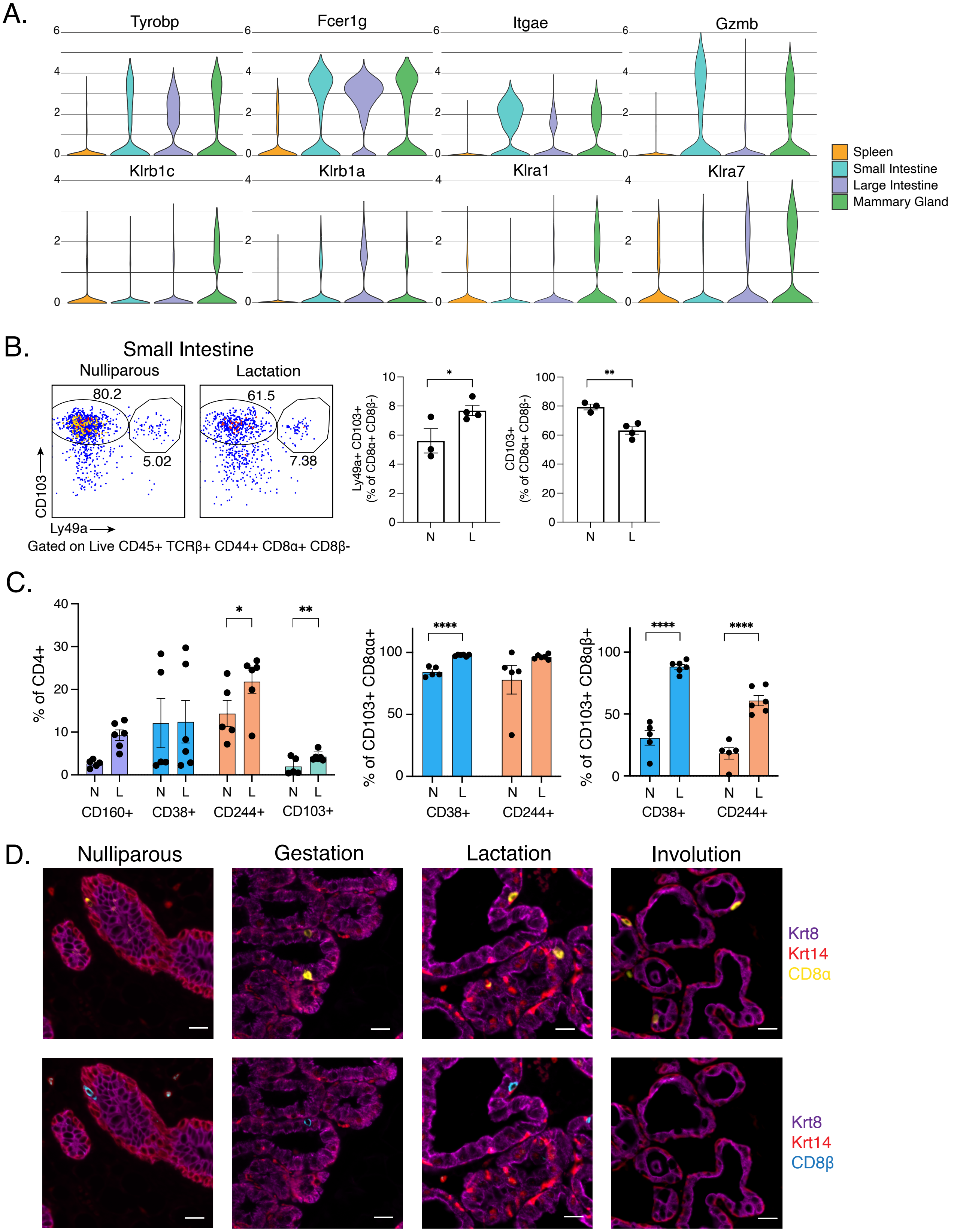

A.
Tyrobp
Fcer1g
Itgae
Gzmb
Klrb1c
Klrb1a
Klra1
Klra7
Spleen
Small Intestine
Large Intestine
Mammary Gland
B.
Small Intestine
Nulliparous
Lactation
80.2
5.02
61.5
7.38
CD103
Ly49a
Gated on Live CD45+ TCRβ+ CD44+ CD8α+ CD8β-
Ly49a+ CD103+ (% of CD8α+ CD8β-)
CD103+ (% of CD8α+ CD8β-)
N
L
C.
% of CD4+
CD160+
CD38+
CD244+
CD103+
% of CD103+ CD8αα+
% of CD103+ CD8αβ+
D.
Nulliparous
Gestation
Lactation
Involution
Krt8
Krt14
CD8α
Krt8
Krt14
CD8β

**Supplementary Figure 5. Mammary T cells display similar markers to intestinal T cells and are intraepithelial in location.**

A) Violin plots depicting gene expression of classical IEL markers in CD8+ T cells from spleen, small intestinal epithelium, large intestinal epithelium and mammary gland of lactating mice.

B) Representative flow cytometry plots and proportion of CD103+ and CD103+ Ly49+ cells in the CD8αα+ population in the small intestine in N=nulliparous (n=3) and L=lactation days 3-5 (n=4).

C) Proportion of CD4+ cells that express CD160, CD38, CD244, and CD103 in N=nulliparous (n=5) and L=lactation days 3-5 (n=6) (left). Proportion of CD103+ cells that are CD38+ and CD244+ in CD8αα+ and CD8αβ+ T cell subsets in nulliparous (n=5) and lactation (n=6) (right).

D) Representative immunofluorescence images of the mammary gland at nulliparous, gestation (G17), lactation (L3), and involution of the epithelium (Krt8, luminal cells, in magenta and Krt14, basal cells, in red), and T cell markers, CD8α (yellow), CD8β (cyan). Selected images are from Figure 2E to show the overlap of cells positive for CD8α and CD8β cells within the epithelial layer. Scale bar = 20μm.

*$p<0.05$, **$p<0.01$, ****$p<0.0001$ by student's t-test. Data representative of ≥3 independent experiments for imaging and flow cytometry, bars in plots indicate mean ± SEM. 2 independent experiments for scRNAseq.

Figure S6

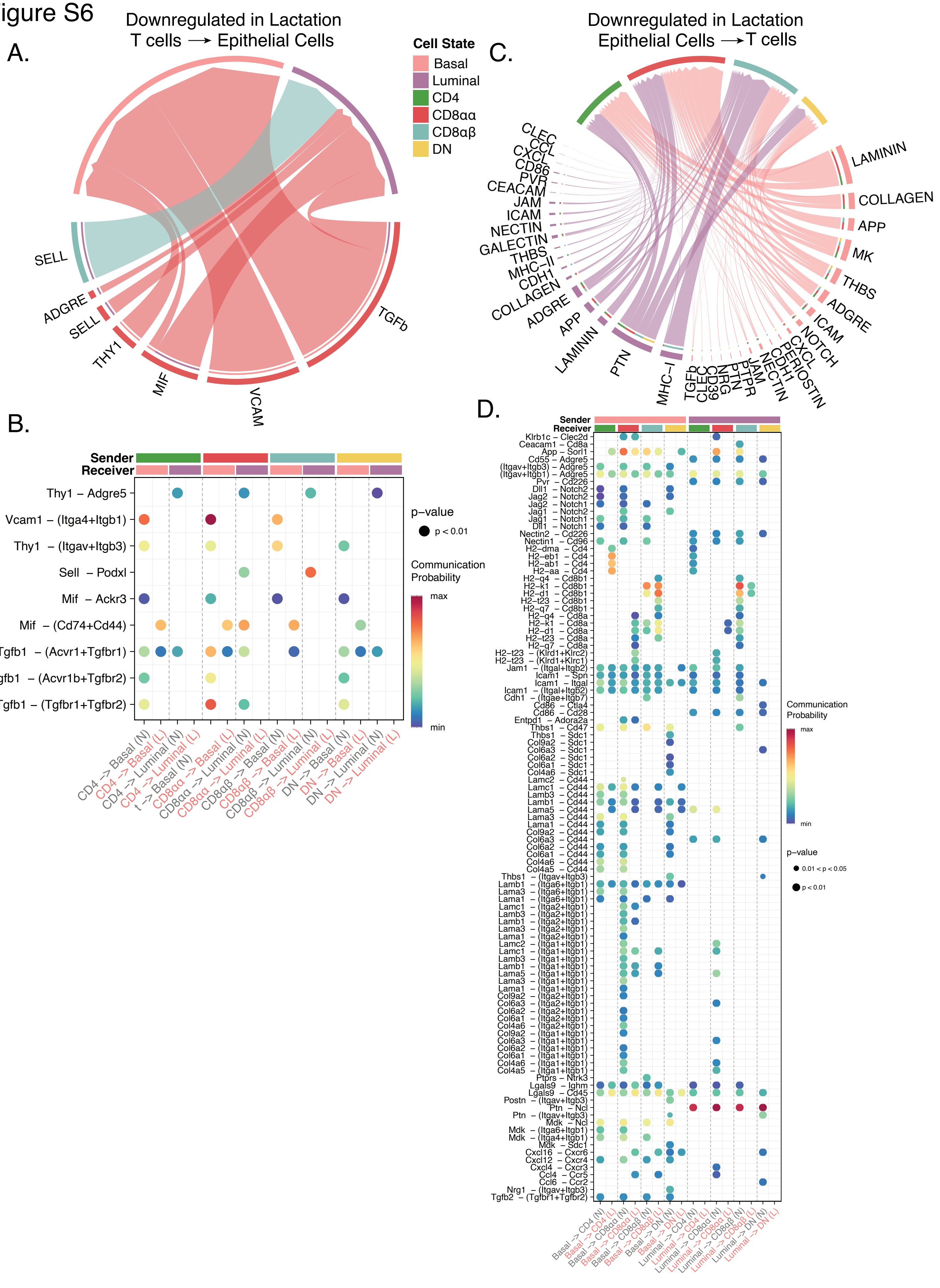

A.
Downregulated in Lactation
T cells ⟶ Epithelial Cells
SELL
ADGRE
SELL
THY1
MIF
VCAM
TGFb
Cell State
Basal
Luminal
CD4
CD8αα
CD8αβ
DN
C.
Downregulated in Lactation
Epithelial Cells ⟶ T cells
CLEC
CCL
CXCL
CD86
PVR
CEACAM
JAM
ICAM
NECTIN
GALECTIN
THBS
MHC-II
CDH1
COLLAGEN
ADGRE
APP
LAMININ
PTN
MHC-I
TGFb
CLEC
CD39
NRG
PTN
PTPR
JAM
NECTIN
CDH1
PERIOSTIN
CXCL
NOTCH
ICAM
ADGRE
THBS
MK
APP
COLLAGEN
LAMININ
B.
Sender
Receiver
Thy1 – Adgre5
Vcam1 – (Itga4+Itgb1)
Thy1 – (Itgav+Itgb3)
Sell – Podxl
Mif – Ackr3
Mif – (Cd74+Cd44)
Tgfb1 – (Acvr1+Tgfbr1)
Tgfb1 – (Acvr1b+Tgfbr2)
Tgfb1 – (Tgfbr1+Tgfbr2)
p-value
p < 0.01
Communication Probability
max
min
CD4 -> Basal (N)
CD4 -> Basal (L)
CD4 -> Luminal (N)
CD4 -> Luminal (L)
CD8αα -> Basal (N)
CD8αα -> Basal (L)
CD8αα -> Luminal (N)
CD8αα -> Luminal (L)
CD8αβ -> Basal (N)
CD8αβ -> Basal (L)
CD8αβ -> Luminal (N)
CD8αβ -> Luminal (L)
DN -> Basal (N)
DN -> Basal (L)
DN -> Luminal (N)
DN -> Luminal (L)
D.
Sender
Receiver
Communication Probability
max
min
p-value
0.01 < p < 0.05
p < 0.01
Basal -> CD4 (N)
Basal -> CD4 (L)
Basal -> CD8αα (N)
Basal -> CD8αα (L)
Basal -> CD8αβ (N)
Basal -> CD8αβ (L)
Basal -> DN (N)
Basal -> DN (L)
Luminal -> CD4 (N)
Luminal -> CD4 (L)
Luminal -> CD8αα (N)
Luminal -> CD8αα (L)
Luminal -> CD8αβ (N)
Luminal -> CD8αβ (L)
Luminal -> DN (N)
Luminal -> DN (L)

**Supplementary Figure 6. Putative interaction networks between mammary T cells and epithelial cells downregulated during lactation.**

A, C) Chord diagrams showing potential signaling pathways downregulated in lactation from T cell populations to epithelial cells (A) and from epithelial cells to T cell populations (C). Ligand:receptor pairs as summarized into functionally related signaling pathways. Outer thicker bars represent the cell population that is the source or target of the signaling pathway in the chord diagram. The inner thinner bar color is the target of the signal. The thickness of the edge represents the signaling strength (communication probability) as calculated by CellChat.

B, D) Dot plots showing the communication probabilities of ligand:receptor pairs downregulated in lactation from T cells to epithelial cells (B) and from epithelial cells to T cell populations (D). Heatmap depicts the communication probability of each ligand:pair for each cell pair in nulliparous (N) and lactating (L) mammary glands. Sender and receivers are indicated by the color bars on top.

### Figure S7

A.

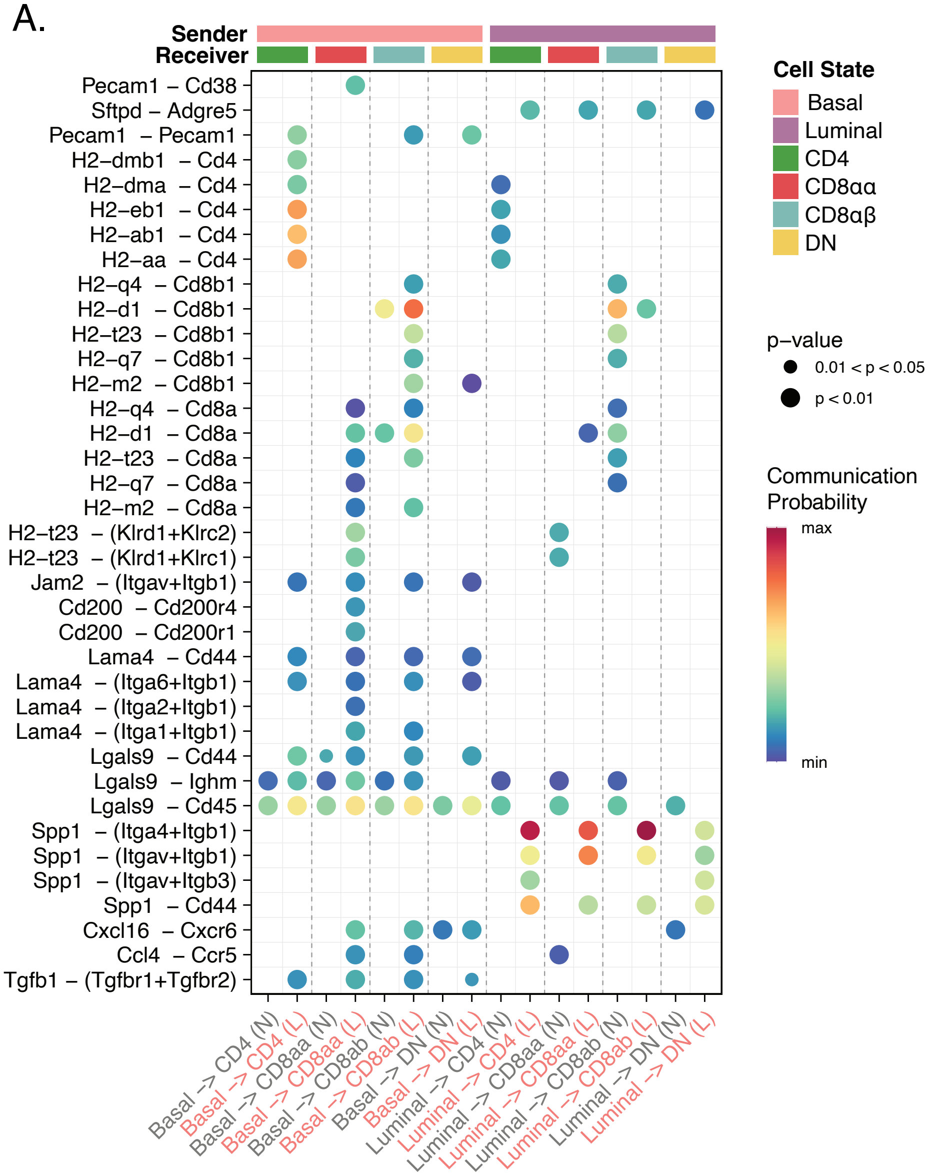

Sender
Receiver
Pecam1 – Cd38
Sftpd – Adgre5
Pecam1 – Pecam1
H2–dmb1 – Cd4
H2–dma – Cd4
H2–eb1 – Cd4
H2–ab1 – Cd4
H2–aa – Cd4
H2–q4 – Cd8b1
H2–d1 – Cd8b1
H2–t23 – Cd8b1
H2–q7 – Cd8b1
H2–m2 – Cd8b1
H2–q4 – Cd8a
H2–d1 – Cd8a
H2–t23 – Cd8a
H2–q7 – Cd8a
H2–m2 – Cd8a
H2–t23 – (Klrd1+Klrc2)
H2–t23 – (Klrd1+Klrc1)
Jam2 – (Itgav+Itgb1)
Cd200 – Cd200r4
Cd200 – Cd200r1
Lama4 – Cd44
Lama4 – (Itga6+Itgb1)
Lama4 – (Itga2+Itgb1)
Lama4 – (Itga1+Itgb1)
Lgals9 – Cd44
Lgals9 – Ighm
Lgals9 – Cd45
Spp1 – (Itga4+Itgb1)
Spp1 – (Itgav+Itgb1)
Spp1 – (Itgav+Itgb3)
Spp1 – Cd44
Cxcl16 – Cxcr6
Ccl4 – Ccr5
Tgfb1 – (Tgfbr1+Tgfbr2)
Basal -> CD4 (N)
Basal -> CD4 (L)
Basal -> CD8aa (N)
Basal -> CD8aa (L)
Basal -> CD8ab (N)
Basal -> CD8ab (L)
Basal -> DN (N)
Basal -> DN (L)
Luminal -> CD4 (N)
Luminal -> CD4 (L)
Luminal -> CD8aa (N)
Luminal -> CD8aa (L)
Luminal -> CD8ab (N)
Luminal -> CD8ab (L)
Luminal -> DN (N)
Luminal -> DN (L)
Cell State
Basal
Luminal
CD4
CD8αα
CD8αβ
DN
p–value
0.01 < p < 0.05
p < 0.01
Communication Probability
max
min

**Supplementary Figure 7. Putative ligand:receptor pairs from epithelial cells to mammary T cells upregulated during lactation.**

A) Dot plot showing the communication probabilities of ligand:receptor pairs upregulated in lactation from epithelial cells to T cell populations (D). Heatmap depicts the communication probability of each ligand:pair for each cell pair in nulliparous (N) and lactating (L) mammary glands. Sender and receivers are indicated by the color bars on top.

Figure S8

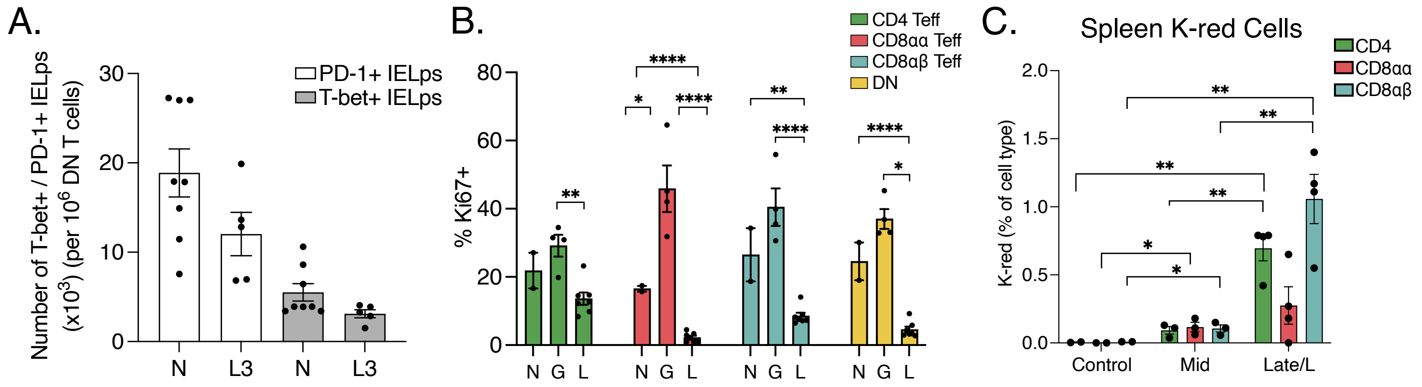

A.
Number of T-bet+ / PD-1+ IELps (x10^3) (per 10^6 DN T cells)
PD-1+ IELps
T-bet+ IELps
N
L3
N
L3
B.
% Ki67+
CD4 Teff
CD8αα Teff
CD8αβ Teff
DN
N G L
C.
Spleen K-red Cells
K-red (% of cell type)
CD4
CD8αα
CD8αβ
Control
Mid
Late/L

**Supplementary Figure 8. Thymic vs intestinal input in mammary T cell expansion.**

A) Representative flow cytometry plots and quantification of total number of T-bet+ and PD-1+ IELps (gated on TCRβ+ CD4- CD8α- CD5+ CD122+ Thy1+) in N=nulliparous (n=8) and L3=lactating day 3 (n=5) thymus.

B) Proportion of T cell populations that are Ki67+. N=nulliparous (n=2), G=gestation day 17 (n=4), and L=lactation days 3-5 (n=7). Teff populations were determined as CD4 Teff: CD4+CD44+CD62L-. CD8αα Teff: CD8α+CD8β-CD44+CD62L-. CD8αβ Teff: CD8α+CD8β+CD44+CD62L-. DN: TCRβ+CD4-CD8α-.

C) Proportion of Kaede red cells within T cell populations (gated on TCRβ+ followed by either CD4+ or CD8β+ or CD8α+ CD8β-) in the spleen 24 hours post-photoconversion of the intestine. Controls are non-photoconverted mice (n=3), mid being mice photoconverted on gestation day 10 and analyzed on gestation day 11 (n=3) and late/L representing mice both photoconverted on gestation day 16 and analyzed on gestation day 17 and mice photoconverted on lactation day 1 and analyzed on lactation day 2 (n=4).

*$p<0.05$, **$p<0.01$, ****$p<0.0001$ by student's t-test. Data representative of ≥3 independent experiments for flow cytometry and Kaede experiments, bars in plots indicate mean ± SEM.

### Figure S9

## A.

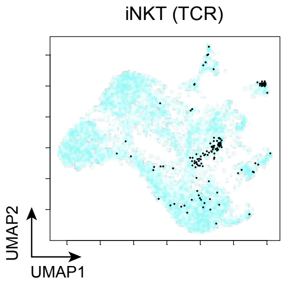


| | alpha.junct | . Cells | Nb Nseq |
|---|---|---|---|
| 3 | CVVGDRGSALGRLHF | 88 | 4 |
| 1 | CVVADRGSALGRLHF | 7 | 3 |
| 5 | CVVVDRGSALGRLHF | 4 | 2 |
| 4 | CVVGDRGTALGRLHF | 1 | 1 |
| 2 | CVVDDRGSALGRLHF | 2 | 1 |

## B.

| | **Mouse** | | **Cell Type** | | | | | | | | | |
|---|---|---|---|---|---|---|---|---|---|---|---|---|
| **Expanded Clonotypes** | LactMG m1 | LactMG m2 | CD4.Tn | CD8.Tn | CD4.Eff | CD8.Eff | CD8aa.Eff | DN | Cycling | Treg | Nb TCRa junct NucSeqs | Nb TCRb junct NucSeqs |
| TRAV6-1.CVLGDLITGNTGKLIF.TRAJ37.TRBV13-2.CASGDWGESQNTLYF.TRBJ2-4 | 16 | 0 | 0 | 0 | 12 | 2 | 0 | 0 | 2 | 0 | 1 | 1 |
| TRAV12D-3.CALVMNNNAGAKLTF.TRAJ39.TRBV12-2.CASSLGLAYEQYF.TRBJ2-7 | 8 | 0 | 0 | 0 | 0 | 1 | 3 | 4 | 0 | 0 | 1 | 1 |
| TRAV4-4/DV10.CAGLNTGNYKYVF.TRAJ40.TRBV14.CASSFRVSNERLFF.TRBJ1-4 | 7 | 0 | 0 | 0 | 5 | 0 | 0 | 0 | 2 | 0 | 1 | 1 |
| TRAV9D-3.CALSAPIDRGSALGRLHF.TRAJ18.TRBV26.CASSRGGGAEQFF.TRBJ2-1 | 0 | 6 | 0 | 0 | 6 | 0 | 0 | 0 | 0 | 0 | 1 | 1 |
| TRAV11.CVVGDRGSALGRLHF.TRAJ18.TRBV13-2.CASGAGGAQDTQYF.TRBJ2-5 | 0 | 6 | 0 | 0 | 0 | 0 | 0 | 0 | 0 | 0 | 1 | 1 |
| TRAV1.CAVRNNYAQGLTF.TRAJ26.TRBV13-3.CASHQGREDTLYF.TRBJ2-4 | 6 | 0 | 0 | 0 | 5 | 0 | 0 | 0 | 1 | 0 | 1 | 1 |
| TRAV8N-2.CATDASSSFSKLVF.TRAJ50.TRBV5.CASSQWGSQNTLYF.TRBJ2-4 | 0 | 5 | 0 | 0 | 3 | 1 | 0 | 0 | 1 | 0 | 1 | 1 |
| TRAV14D-1.CAATLSGSFNKLTF.TRAJ4.TRBV13-3.CASSPQEGYAEQFF.TRBJ2-1 | 4 | 0 | 0 | 0 | 3 | 0 | 0 | 1 | 0 | 0 | 1 | 1 |
| TRAV13D-2.CAIDPNYNQGKLIF.TRAJ23.TRBV31.CAWSLGGFYAEQFF.TRBJ2-1 | 0 | 5 | 0 | 0 | 3 | 1 | 1 | 0 | 0 | 0 | 1 | 1 |
| TRAV13-2.CAIGGTGNTGKLIF.TRAJ37.TRBV26.CASSLSRNSGNTLYF.TRBJ1-3 | 5 | 0 | 0 | 0 | 5 | 0 | 0 | 0 | 0 | 0 | 1 | 1 |
| TRAV9-2.CVFPQNYNQGKLIF.TRAJ23.TRBV26.CASRGQNYEQYF.TRBJ2-7 | 0 | 4 | 0 | 0 | 3 | 0 | 0 | 0 | 1 | 0 | 1 | 1 |
| TRAV7D-5.CAVSSNYNQGKLIF.TRAJ23.TRBV3.CASSAGTGGIEQYF.TRBJ2-7 | 0 | 4 | 0 | 0 | 0 | 0 | 0 | 0 | 4 | 0 | 1 | 1 |
| TRAV7D-4.CAASEANSGTYQRF.TRAJ13.TRBV26.CASSRGGGAEQFF.TRBJ2-1 | 0 | 4 | 0 | 0 | 2 | 0 | 0 | 0 | 2 | 0 | 1 | 1 |
| Nb of cells | 2242 | 3072 | 835 | 797 | 536 | 1461 | 857 | 275 | 191 | 87 | | |

**Supplementary Figure 9. TCR clonotypes expand across different T cell types in the lactating mammary gland.**

A) UMAP and quantification of cells with different alpha junction usage.

B) Quantification of expanded clonotypes across mice and cell types.

### Figure S10

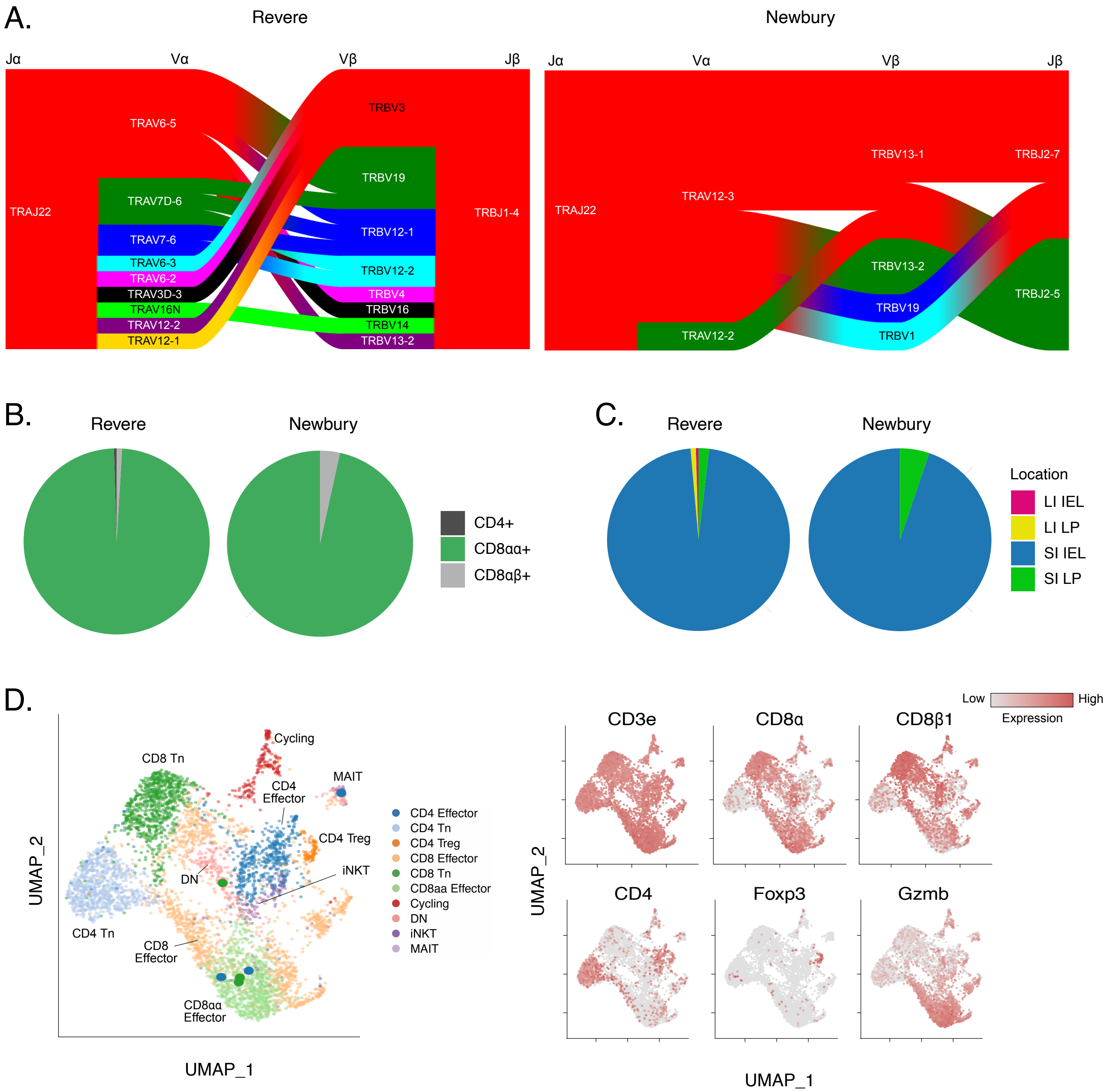
A.
Revere
Jα
Vα
Vβ
Jβ
TRAJ22
TRAV6-5
TRAV7D-6
TRAV7-6
TRAV6-3
TRAV6-2
TRAV3D-3
TRAV16N
TRAV12-2
TRAV12-1
TRBV3
TRBV19
TRBV12-1
TRBV12-2
TRBV4
TRBV16
TRBV14
TRBV13-2
TRBJ1-4
Newbury
Jα
Vα
Vβ
Jβ
TRAJ22
TRAV12-3
TRAV12-2
TRBV13-1
TRBV13-2
TRBV19
TRBV1
TRBJ2-7
TRBJ2-5
B.
Revere
Newbury
CD4+
CD8αα+
CD8αβ+
C.
Revere
Newbury
Location
LI IEL
LI LP
SI IEL
SI LP
D.
Cycling
CD8 Tn
MAIT
CD4 Effector
CD4 Treg
iNKT
DN
CD4 Tn
CD8 Effector
CD8aa Effector
UMAP_2
UMAP_1
CD4 Effector
CD4 Tn
CD4 Treg
CD8 Effector
CD8 Tn
CD8aa Effector
Cycling
DN
iNKT
MAIT
Low
High
Expression
CD3e
CD8α
CD8β1
CD4
Foxp3
Gzmb
UMAP_2
UMAP_1

**Supplementary Figure 10. Revere and Newbury TCRs have distinct V region unsafe and are only expressed in CD8αα+ T cells in the small intestine and mammary gland.**

A) V region usage of Revere and Newbury TCR families.

B) Quantification of Revere and Newbury families in different T cell types in the small intestine.

C) Quantification of Revere and Newbury families in the IEL and lamina propria compartments in the small intestine and large intestine.

D) UMAP projection of lactating mammary gland T cells with highlighted dots representing cells expressing Revere and Newbury TCRs (left) with feature plots of T cell genes (right).

Data representative of 2 independent experiments.

Figure S11

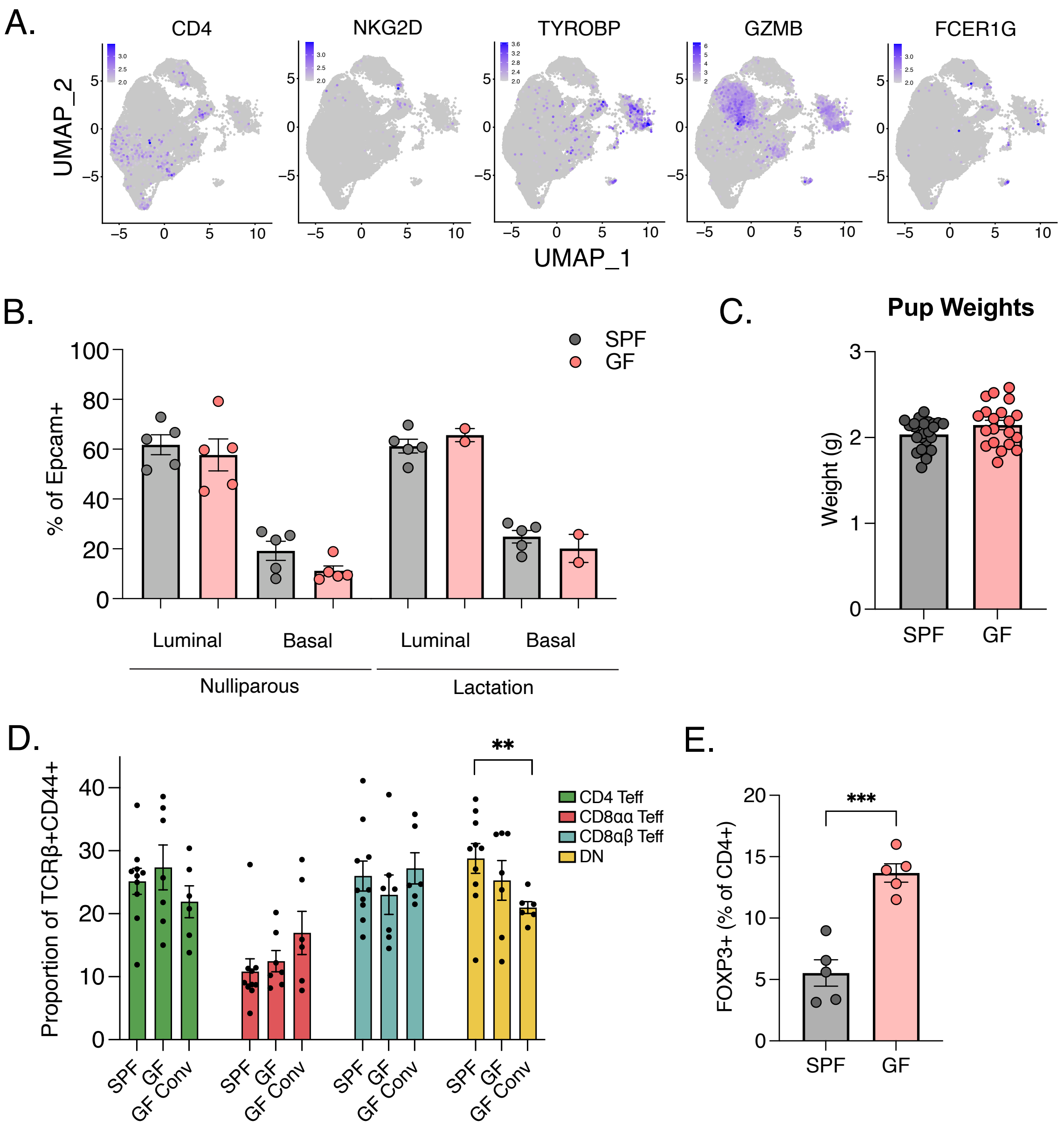
A.
CD4
NKG2D
TYROBP
GZMB
FCER1G
UMAP_2
UMAP_1
B.
SPF
GF
% of Epcam+
Luminal
Basal
Luminal
Basal
Nulliparous
Lactation
C.
Pup Weights
Weight (g)
SPF
GF
D.
Proportion of TCRβ+CD44+
CD4 Teff
CD8αα Teff
CD8αβ Teff
DN
**
SPF
GF
GF Conv
E.
FOXP3+ (% of CD4+)
***
SPF
GF

**Supplementary Figure 11. Characterization of human IEL-like cells and lactogenesis in germ-free mice.**

A) Feature plots of select CD8αα+ Teff markers, cytotoxic markers and tissue resident genes projected on UMAP from human breast tissue (6A).

B) Proportion of basal (sort marker) and luminal (sort marker) Epcam+ cells in SPF (n=5) and GF (n=5 nulliparous and n=2 lactating) nulliparous and lactating mammary glands.

C) The weight (grams) of pups from GF (n= 3 litters) and SPF (n= 4 litters) litters at 4 days of birth normalized to litter size.

D) T cell population proportions of TCRβ+ CD44+ cells of SPF (n=10), GF (n=7), and GF conventionalized (n=6) mammary glands. Teff populations were determined as CD4 Teff: CD4+CD44+CD62L-. CD8αα Teff: CD8α+CD8β-CD44+CD62L-. CD8αβ Teff: CD8α+CD8β+CD44+CD62L-. DN: TCRβ+CD4-CD8α-.

E) Proportion of mammary Foxp3+ T regulatory cells in lactating SPF (n=5) and GF mice (n=5).

**$p<0.01$, ***$p<0.001$ by student's t-test. Data representative of ≥ 3 independent experiments, bars in plots indicate mean ± SEM.
